# GlyNet: A Multi-Task Neural Network for Predicting Protein-Glycan Interactions

**DOI:** 10.1101/2021.05.28.446094

**Authors:** Eric J. Carpenter, Shaurya Seth, Noel Yue, Russell Greiner, Ratmir Derda

## Abstract

Advances in diagnostics, therapeutics, vaccines, transfusion, and organ transplantation build on a fundamental understanding of glycan-protein interactions. To aid this, we developed GlyNet, a model that accurately predicts interactions (relative binding strengths) between mammalian glycans and 352 glycan-binding proteins, many at multiple concentrations. For each glycan input, our model produces 1257 outputs, each representing the relative interaction strength between the input glycan and a particular protein sample. GlyNet learns these continuous values using relative fluorescence units (RFUs) measured on 599 glycans in the Consortium for Functional Glycomics glycan arrays and extrapolates these to RFUs from additional, untested glycans. GlyNet’s output of continuous values provides more detailed results than classification models. Such continuous outputs are easily converted by a following classifier, and in this form GlyNet outperforms reported classifiers. GlyNet is the first multi-output regression model for protein-glycan interactions and will serve as an important benchmark, facilitating development of quantitative computational glycobiology.

---

A complex array of carbohydrates (glycans) coat the surfaces of every cell^1^. Interaction of this glycan coat with protein receptors on human cells is the first step in distinguishing human cells from invading pathogenic bacteria or viruses, or detecting cancerous cells in human tissues^2^. A major tool for collecting glycomic data, the glycan array, is akin to DNA microarrays:^3^ a variety of carbohydrates are deposited onto distinct locations on a glass surface^4^ or distinct beads. Such arrays can simultaneously measure the binding strengths of several hundred carbohydrates with a given sample of a glycan binding protein^5,6^ (GBP), producing a glycan binding profile for that protein. This data is a critical starting point for fundamental applications such as improved designs of inhibitors, vaccines and therapeutics^7–12^. Despite the fundamental importance of glycans, their investigation is orders of magnitude slower than the study of proteins or nucleic acids, for several reasons: (i) Glycans are not directly encoded by DNA^1^: thus, “glycomics” information cannot be collecting using powerful next-generation sequencing techniques. (ii) As glycans are not linear but branched structures, analytic techniques for linear sequences cannot be readily applied to their graph structures. (iii) Laboratory synthesis of glycan oligomers is significantly more difficult than the synthesis of amino acid oligomers (proteins) and RNA/DNA oligomers. Predicting the strength of the interaction between any given glycan and protein requires methods that can effectively extrapolate beyond our limited experimental knowledge of glycomics.

The ability to predict protein-glycan interactions, even approximately, reduces the search space that needs to be explored by experiments. Not only is the space of glycans difficult to explore, it also scales significantly faster than the analogous space of DNA-RNA or proteins. For example, considering glycans composed of just 10 common mammalian monosaccharides with 2×4 different linkages between them, {α, β} × {1-2, 1-3, 1-4, 1-6}, we count 56,880 trisaccharides. It is not currently feasible to synthesize this set, much less explore their interactions. These are small structures, our glycan count increases to 4.4×10^6^ tetrasaccharides and 3.8×10^8^ pentasaccharides (see Supplementary Table S1 for a list of these structures); enlarging the constituents to the dozens of monosaccharides and considering additional linkage variety, the number of possible glycans grows even larger. Collective human knowledge in glycomics falls several orders of magnitude short. For example, GlyTouCan 3.0, today’s most exhaustive database of glycan properties, contains information about only 120,000 entries containing more than 100 diverse monosaccharides.^13^ One critical property, is whether a specific glycan interacts with a particular protein, and how strongly; datasets that describe glycan-protein interactions contain only a small subset of those glycan structures^14^. Advances in understanding peptide and nucleic acid sequences is fueled by tools that give rise to experimental libraries of millions of instances and high-throughput sequencing that can exhaustively analyse them. Unfortunately, state-of-the-art glycan synthesis approaches generate libraries of only hundreds of glycans^15,16^; and parallel testing of glycan properties on glycan arrays yields information about hundreds of structures. The fundamental challenges in synthesizing glycans and collecting new glycomics data motivated our work to extrapolate beyond the existing glycan-protein binding data.

Previous machine learning (ML) approaches to protein-glycan interaction employed techniques like support vector machines^17,18^, graph kernels^18^, modularity optimization methods^19^, and Markov models^20–22^, to identify glycan motifs, substructures that specific proteins recognize–for reviews see Mamitsuka^23^, Haab^24^, and Sese^25^. Several publications from Aoki-Kinoshita and co-workers^20,22,26–29^, Coff and co-workers^30^, Cummings and co-workers^14^, and recently Bojar and co-workers^31–34^ focus on ML *classification models* that predict qualitative features (think “strong *vs.* weak” interactions) for each glycan-protein pair. Woods and co-workers combined molecular mechanics, automated 3D glycan structure generation and docking techniques to produce computational carbohydrate grafting that can qualitatively predict binding between a carbohydrate fragment and a known 3D protein structure^35^. Malik and Ahmad provide a learned model for the reverse task, asking whether a protein has (learned) features, to predict whether it binds to a particular glycan^36^.

In recent years, ML algorithms based on Neural Networks (NN)^37,38^ have achieved remarkable success classifying images, sound, text and linear biological oligomers (protein, DNA, RNA). Dahl^39^, Ma^40^, and Jensen et al.^41^ pioneered the use of deep NN to predict physicochemical properties of organic molecules. Neural networks have also been used to predict patterns of enzymatic processing on glycopeptides^38^, taxonomic classifications of organisms synthesizing glycans^33^, and biological properties like immunogenicity of glycans^32,34^. At the time of the preparation of this manuscript, Bojar and coworkers released a bioRxiv manuscript describing the first graph neural network (GNN) that predicts properties of glycan graphs such as evolutionary origin, immunogenicity and recognition by viral proteins^34^. Here we attempt to advance the state-of-the-art by developing a regression model with *quantitative* outputs of protein-glycan interaction using neural networks.

Binding of a given glycan to a specific protein is fundamental to biology, and the strength of this interaction—aka affinity and avidity—are the foundation of most biological responses elicited by glycans. Here, we develop a NN that takes a glycan structure as an input and outputs quantitative binding values, i.e. the “strength” of the interaction, for specific proteins. Training a NN typically requires large numbers of labelled instances–here, sets of protein-glycan pairs, each labeled with the associated K_d_ (avidity and affinity) values. Unfortunately, there are no such datasets readily available. However, as semi-quantitative estimates of glycan-protein binding strength we use relative fluorescence unit (RFU) values obtained in glycan array experiments conducted by the Consortium for Functional Glycomics (CFG)^4^. In this manuscript, we propose a system in which a glycan is encoded as an input into a neural network, which then predicts the RFU values for that glycan across a large set of protein samples (Fig. 1). We use 752,943 RFU values from the binding of 599 glycans to 353 proteins at different concentrations (1257 samples total) to train the neural network, providing the first example of a regression model which predicts continuous RFU values for an input glycan across this diverse set of protein samples; here based on a multi-output neural network architecture.

**Fig 1.**
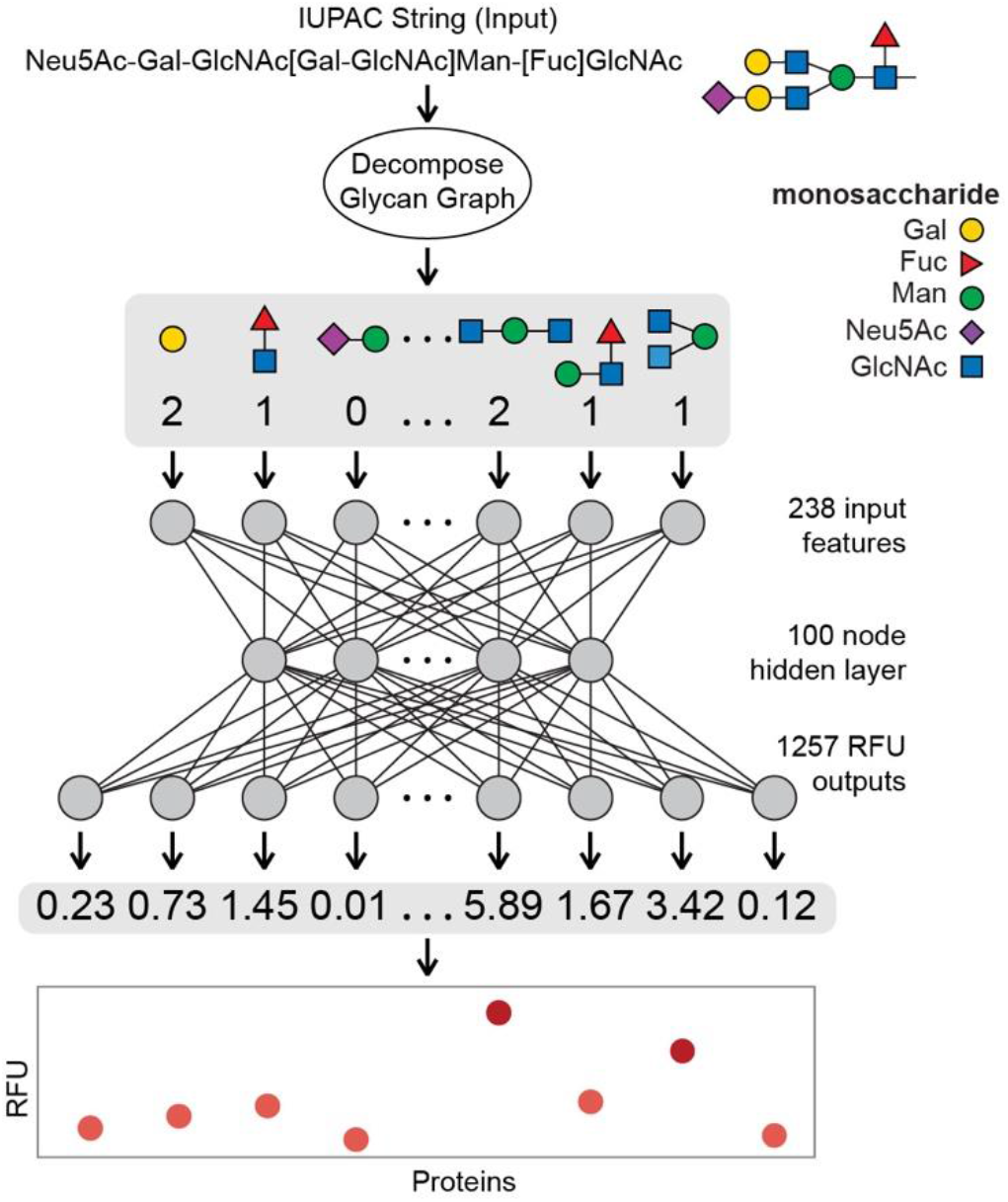
Schematic overview of the work described in this paper. Each glycan is decomposed into a fingerprint of feature counts, primarily of small *q*-gram subtrees. This fingerprint is then input into the learned model which outputs the predicted binding value between the glycan and 1257 proteins. Once trained, the networks can be used to predict binding of novel glycans to the proteins.

## Results

### Selection of CFG Glycan Array Protein Binding Data

In general, a glycan array contains several dozen to several hundred glycans, each printed on one or more discrete spots; when exposed to a given protein sample, one can then read off the RFU for each spot. Combining the data from over a thousand such experiments provide convenient and uniform data for training and large-scale data analysis.^19,25,28,30^ Specifically, we combine data from CFG’s Mammalian Printed Array v5.0, v5.1 and v5.2 arrays, taking the mean of the six replicates. We omit the two glycans not available on the v5.2 arrays, and two glycans with errors in the reported structure. We also omitted eight glycans containing one of five rare monosaccharides (each appearing four or fewer times among all the glycans: Rha, GlcN(Gc), G-ol, MurNAc, Neu5,9Ac2). Specifics of all the omissions are available in the Supplementary Information. These glycans range from 1 to 36 monosaccharides in size (mean = 6.1, median = 4, Fig. 2A), they collectively contain 10 different monomers and 12 anomeric linkages (Fig. 2B). Supplementary Table S2 lists the 599 glycans and the omitted ones. We omitted glycan data prior to ver. 5.0 CFG arrays because these arrays contain fewer glycans then Ver 5.0-5.2 arrays. Merger of this data with Ver 5.0-5.2 arrays would create an unbalanced dataset in which data describing interactions of some glycans with some protein samples is non-existent and datasets with missing data are less convenient for training than the complete datasets (Fig. 3A).

**Fig. 2.**
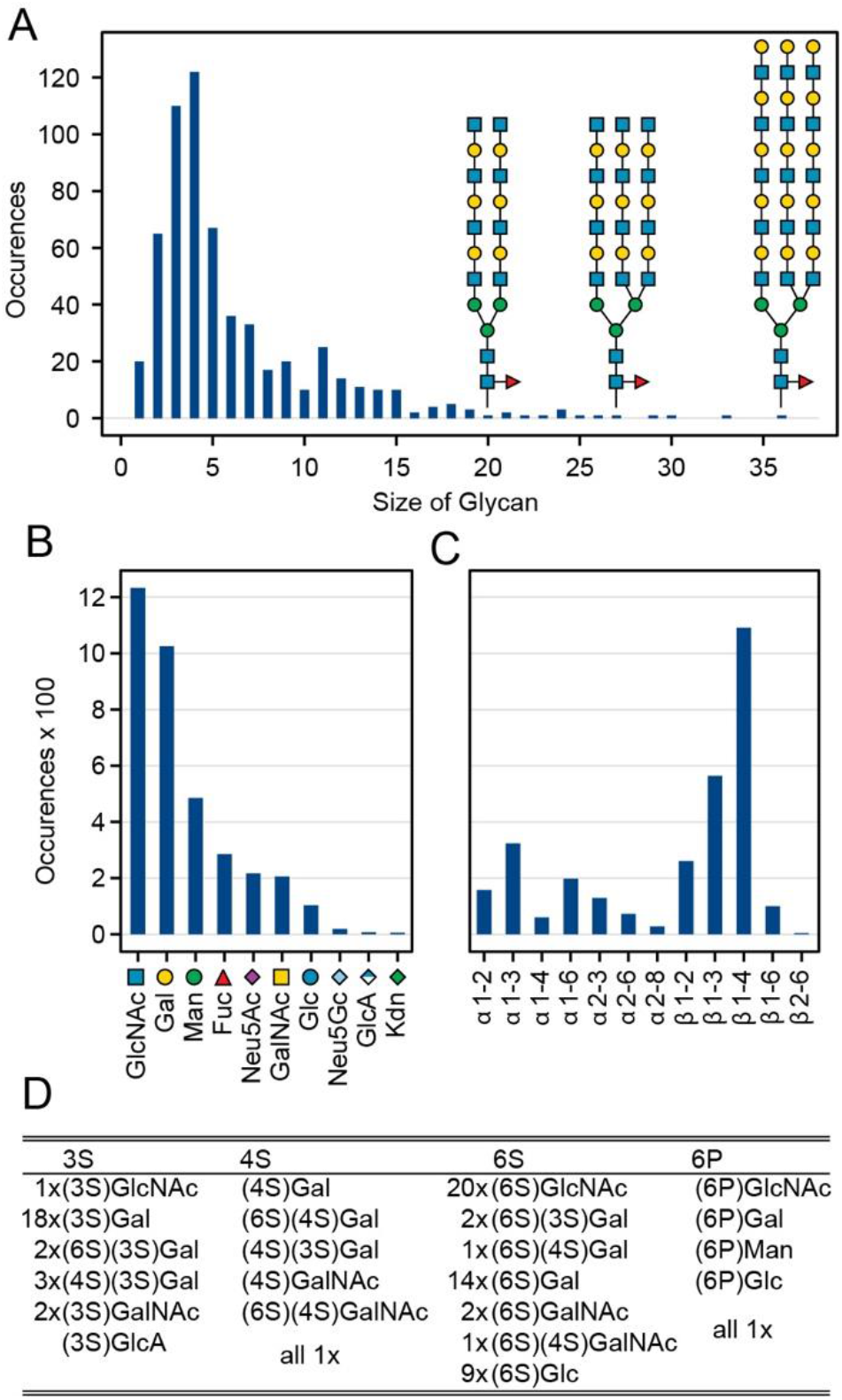
Glycans used for training of GlyNet. A) Distribution of the glycan sizes and examples of bi- and triantennary glycans with 20, 27 and 36 monosaccharides. B) Distribution of the monosaccharides within the 599 glycans. C) Distribution of the different types of the anomeric links within the glycans. D) Details of the modified monosaccharides, showing for each of the four modifications/positions a list of the monosaccharides with that modification and their frequencies.

**Fig. 3.**
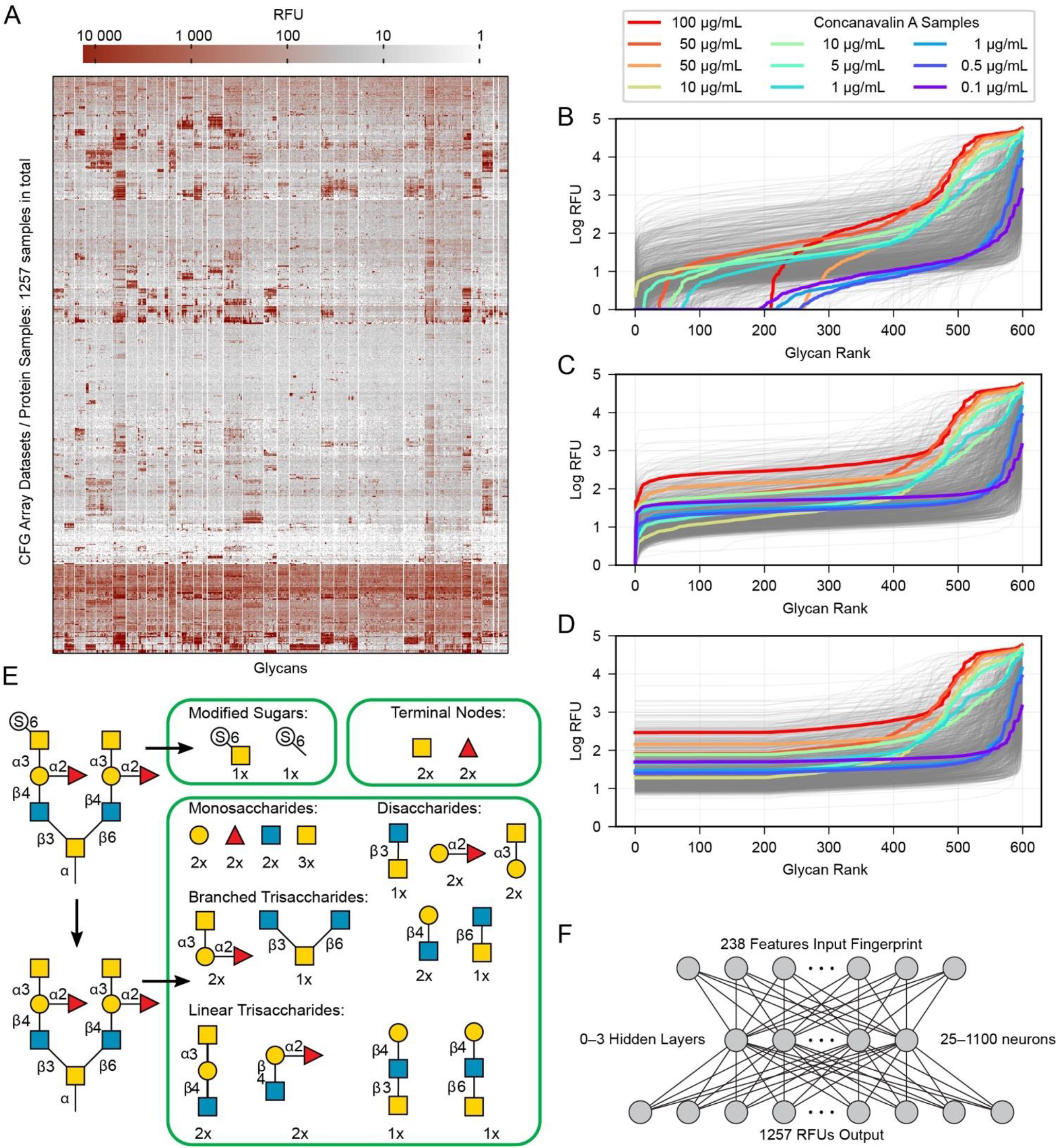
Pre-processing of data. A) Heatmap of the data overall, with all glycans and protein samples. B) The 599 RFU values for each protein are sorted, log-transformed and plotted as a line. Highlighted in color are several samples of the protein Concanavalin A at various concentrations. (Note that due to the sorting, glycans appear at different *x*-axis positions in each line.) C) The raw values are shifted to place the minimum at 1.0. D) The lowest 1/3^rd^ (200 values) are rounded up to the 200^th^ value to remove noise from the training data. E) Details of the subgraph features used to describe each glycan. F) Diagram illustrating the neural network architectures we used in this work. One hidden layer is shown, although we tried zero to three hidden layers.

In addition to ten main monosaccharides, there are phosphorylated or sulfurated units, listed in Fig. 2D. We extracted raw data from a table in the ImaGene^42^ program’s format and chose the 1257 glycan array experiments for which this table is available, see Supplementary Tables S2 and S3 for a full list of the protein samples and glycans including their associated CFG labels. Fig. 3A shows all of the data as a heatmap, similar to analyses by Sese,^25^ a larger version of this panel with labels is available as Supplementary Fig. S1. Overall, 881 of the protein samples correspond to 352 unique proteins (cpbIds); the remaining 350 are complex samples (serum, etc.). Many of the protein samples were tested at several different concentrations (colored lines highlight Concanavalin A in Fig. 3B-D). Specifically, binding of 18 proteins have been measured at 5 or more different concentrations and 69 proteins were measured at 4 distinct concentration (Supplementary Table S3). Since the concentration scan data are relatively rare in the CFG dataset and concentrations are not available for ~120 samples, we treat each protein-concentration pair as a unique sample and consider its binding with respect to the 599 glycans.

### Preprocessing of glycan-protein binding data

For each glycan-protein pair, the glycan arrays measure Relative Fluorescence Units (RFUs) which are a measure of the binding strength for that pair. As each glycan occurs six times in the array, we use the mean over these six replicate array spots for each glycan. We modify these input values in three steps: (i) we add a constant to each RFU values for each glycan array to set the minimum value in each dataset to 1.0 (Fig. 3B) and (ii) log-transformed the values (Fig. 3C) to reduce the several orders-of-magnitude of dynamic range (similar to a previous report^30^). (iii) We then replaced the lowest 1/3^rd^ of these values (data for 200 of 599 glycans) with the value equal to that of the 200^th^ lowest point in that dataset (Fig. 3D). This transformation eliminates the noise/variability at the lower end of the log-transformed dataset caused by fluctuations in the readout machinery, wash conditions and other factors not relevant to the task of learning glycan-protein interactions. Note that the weakest RFUs are non-binders, and we observed that the lowest third contains the worst noise. Filtering more risks removing glycan-protein binding data. The filtering simplifies the learning task, as the model learned from the training process is not trying to reproduce any of the irrelevant patterns in this part of the data. The pre-processed log-mean RFU values are available in Supplementary Table S4.

We selected this method of preprocessing of the CFG data based on a combination of that used in a previous report^30^ and general knowledge about the nature of the RFU data. Whether other approaches (including omitting pre-processing) could yield better results is still an open question. Ideally, various pre-processing approaches should be tested as part of the overall optimization of the learning model (e.g., data from Fig. 3B, 3C, 3D as well as data without any pre-processing could be used as separate instances of ground truth and accuracy of the models trained using these datasets can be compared to one another); however, we do not do such analysis in this manuscript.

### Representation of Glycans for Input to the Neural Networks

In order to input a glycan structure into the neural network, we adapted the *q*-gram^43^ approach, encoding each glycan as a “fingerprint”, i.e., a feature vector containing counts of how often each feature occurs. The major features we included were the contiguous 1, 2, and 3-monosaccharide subgraphs of the glycan structure including the connecting anomeric linkages (Fig. 3E). Including the smaller subgraphs allows the 85 mono- and di-saccharides to be represented (14 % of the glycans), as they have no tri-saccharide subgraphs. Counts of the terminal monosaccharides, the most exposed in protein-glycan interactions, were also used as part of the feature set. Phosphorylation and sulfation of glycans was represented by adding a feature for the phosphate or sulfate group position, a feature for each such modified monosaccharide, and by using the unmodified sugar in the structure used for subgraph generation and terminal positions. This encoding helps to avoid a combinatorial explosion in the subgraphs created by these variants. The feature vector for each glycan contains one element for each feature (272) found in any of the 599 glycans. The corresponding training set is thus a 272 features x 599 glycans table, which is available in Supplementary Table S5. Each of the 599 glycans in the CFG set we selected has an associated set of 1257 RFU values for each protein.

This fingerprint encoding does not describe every feature of the glycan, in particular it does not encode the location of a feature, only how often it occurs. Two (or more) distinct glycans may have identical feature vectors and are thus treated as identical. In the set of 599 glycans, we have three feature vectors encoding different glycan structures (Supplementary Fig. S2). Also for 105 of the 599 glycan instances the CFG description does not record whether the anchoring monosaccharide is in the alpha or beta conformation (Supplementary Table S2). To avoid this ambiguity, we ignore the stereochemistry of the anomeric carbon of this position and similarly, we do not encode which of the CFG spacers (Supplementary Table S6) links the glycan to the substrate. The consequence of these design decisions is an inability of the network to distinguish glycans that may exhibit different binding properties. For example the binding of Gal(β1-3)GalNAc to Fm1D has more than two-orders of magnitude difference across the selection of α-Sp8, α-Sp14, α-Sp16, and β-Sp8, the structures and stereochemistry of the anomeric linkers present in the CFG data (Supplementary Table S2). When we train the network, all four of these are used as distinct input instances and the model output is evaluated with respect to each of the corresponding outputs. But the model can produce only one prediction for Gal(β1-3)GalNAc binding to Fm1D. As the model produces the same output for all these indistinguishable inputs, the prediction can never exactly match, but will be optimized towards a compromise, in our case the means of the CFG values is optimal. However, these instances are not conflated with the Gal(**α**1-3)GalNAc ones because the stereochemistry of the other subunits is input to the network. Our encoding assigns 599 CFG glycans to 515 distinguishable cases, while other encodings might avoid such indistinguishability of glycans, the neural network may not be able to extract patterns of glycan-protein binding from them. Despite the apparent limitation of indistinguishable glycans, we believe our system achieves sufficient accuracy to be useful.

Another issue that creates indistinguishable glycans is that the fingerprint encoding does not encode the location of a feature, only how often it occurs. It is thus possible that two (or more) distinct glycans may have identical feature vectors and are thus treated as identical. From 599 glycans, we found only three examples of different glycans encoded by the same feature vector (Supplementary Fig. S2). These glycans are used as distinct inputs and the model output is evaluated with respect to the corresponding output and the model is optimized to match the mean of the CFG values from two different glycans.

### Multi-task Network

We learned the final multi-task model, named GlyNet, on all the CFG data. To find the best model the learner used *k* = 10-fold internal cross-validation (CV) to identify the appropriate number of hidden layers, the number of nodes in each hidden layer, as well as hyperparameter settings (Fig. 4; RFU predictions used are available in Supplementary Table S7). We randomly partitioned the glycan samples into ten folds, trained a model using the glycans in nine of the folds and validated it on the tenth fold with MSE as the loss function. We ensured that glycans encoded by the same fingerprints (see above) are partitioned to that same fold. By repeating this process for all ten folds with separate models, we obtained the mean performance of our architecture on the data. Optimization used ADAM^44^ implemented in the PyTorch^25^ ML library with batches of 64 instances; we stopped training when the Mean Squared Error (MSE) did not improve for 10 epochs.

**Fig. 4.**
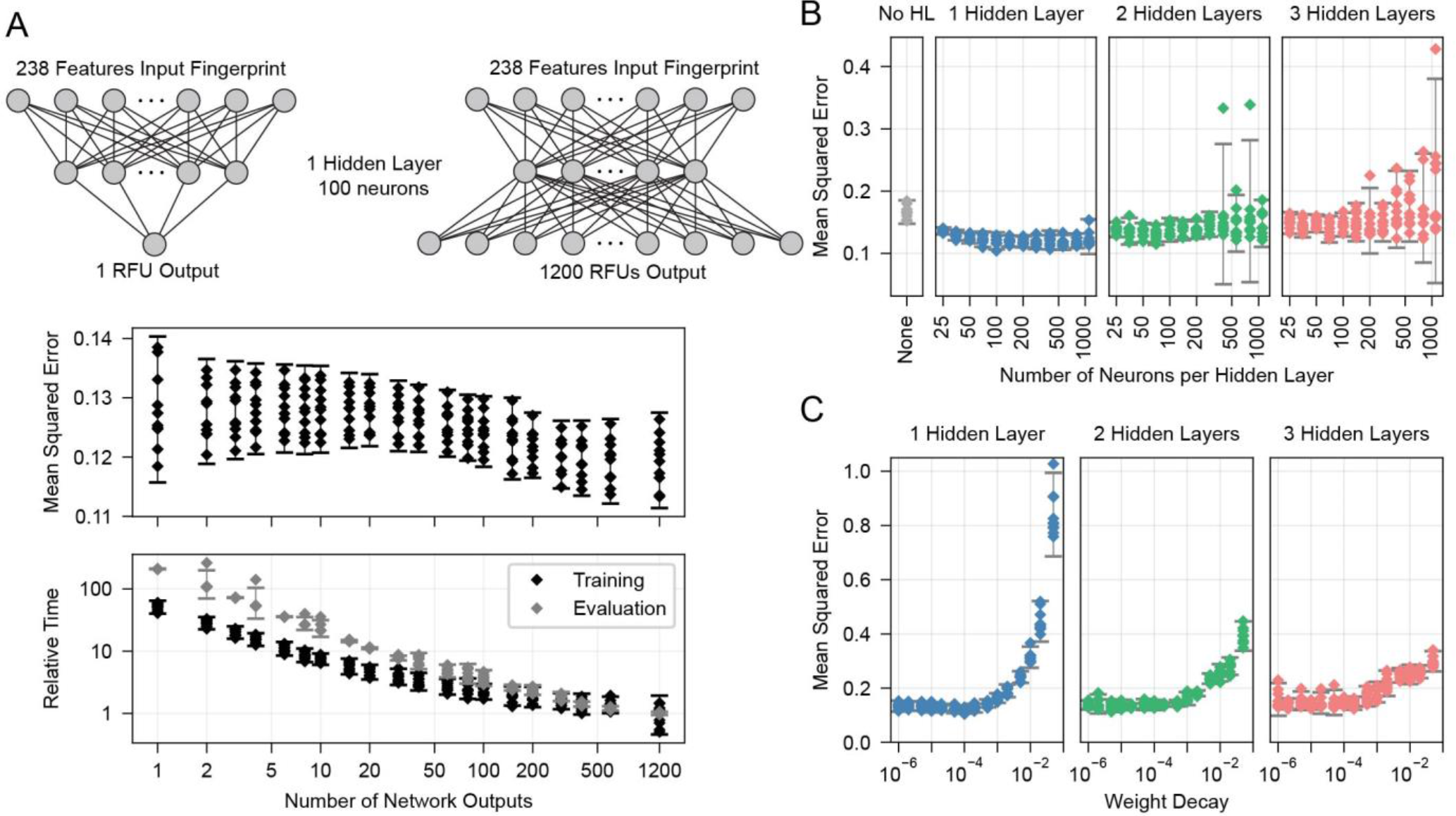
Searches were used to find optimal parameters and settings. A) Effect of varying the number outputs of the neural networks across a subset of 1200 protein samples. Upper panel, the effect on the mean-squared error. Lower panel, relative training and evaluation times. Evaluation time points are mostly consistent and overlap except for occasional outliers. B) The effects of altering the number of hidden layers and their number of neurons. C) Effect of varying the weight decay parameter of the ADAM optimiser. Each error bar is from *n* = 10 points, one from each of the 10 folds of a single run across 599 glycans. Error bars show the 95% confidence interval for a Gaussian of the mean and standard deviation of the plotted points.

Models that predict protein-glycan interactions could be constructed as several separate tasks, with one learned model for producing each RFU prediction. The multi-task approach, adapted from Dahl, et al.^39,40^ was an important development as compared to the single-task approach. Simultaneously outputting data for all protein samples from one model instead of training a separate model for each, greatly reduced overall training time and allowed for a more extensive hyperparameter search. We found that increasing the number of output neurons improved the performance of the model (Fig. 4A). The multi-output network has a better mean squared error (over all 599 training glycans) than a single output network on 75% of the protein/concentration pairs, with a median improvement in the MSE of 10%. We thus focused on one model which given an input glycan simultaneously outputs all 1257 (logarithmically transformed) RFU predictions. As shown in Fig. 3F, our neural network architecture first translates the input glycan into intermediate “hidden layers”, which are then used to generate the (log scaled) RFU values for the 1257 protein-concentration instances. For this to work, the hidden layers need to implicitly represent the glycan characteristics that determine the RFU values for all the proteins. Note that each training instance (one glycan and 1257 RFUs) provides lots of feedback (1257 values) for each of the “intermediate” weights, which suggests it may be relatively easy to train this structure compared to the single output approach. Further, from general knowledge of protein-glycan interaction; many proteins share a set of common rules and there is an expected pattern for the RFU values across the proteins.^45^ In experiments with intermediate cases, multi-output networks producing outputs for only some of the protein samples, there was a minimal size of ~10 outputs below which no improvement was seen, with gradual improvements as the number of outputs increase (Fig. 4A). Also convenient was the decrease in training and evaluation times with greater numbers of outputs (Fig. 4A).

### Network Training

We found a single hidden-layer of 100 nodes optimized with weight decay of 10^−4^ to be the best. In a similar learning task, Dahl^39^ and Ma *et al*^40^ previously reported that multilayer architecture could yield a better performance than a single layer. It is possible that with more data, a deeper architecture could yield a better performance; however, given a rather limited dataset, we opted to proceed with simpler single-layer architecture.

We used neurons with biased ReLU activations for all layers except the final layer which uses biased linear activations instead. Initial versions of the multi-task networks with ReLU in the final layer failed to train properly, with a small number (~20) of RFU outputs producing zeros for all input glycans. These outputs had become trapped in a zero-gradient state and were not being optimized in the learning process. This problem was avoided with linear output neurons which have a non-zero gradient everywhere.

### Evaluation of the Glycan Binding Model by Glycan

After selecting the optimal architecture and producing models on the 9⁄10^ths^ of the training data we compared outputs on the held-out tenth folds with true RFU data obtained from the CFG arrays. Over all 599 glycans for each of the 1257 proteins we achieved a cross-validation mean-squared error (MSE) of 0.12 and mean R^2^ = 0.78 (coefficient of determination – a measure of correlation) in predicting logarithmically transformed RFU values.

For each glycan input to our model, RFU predictions are output for each of the 1257 protein samples (available in Supplementary Table S8) can be compared against the reference CFG measurements to yield MSE for individual glycans. They vary by more than an order of magnitude from 0.036 to 0.495 and Fig. 5 compares prediction for glycans with lowest MSE = 0.036 (Fig. 5B) a median MSE (Fig. 5C) and the highest MSE=0.495 (Fig. 5D). Predictions are denoted with dots and the ground truth, log(mean RFU), is depicted as solid line with a 95% confidence interval (gray band, calculated from six experimental replicates of the RFU data). While we do not use the confidence interval, neither for training, nor for calculation of MSE, we noted that many predictions reside within the experimentally determined confidence interval of RFU values. In other words, performance of the model, while not perfect, is comparable to the experimental variance of the data. Each plot has a list of the protein samples with the highest 20 predicted RFU values. These top-20 predictions further highlight that the model predictions accurately reproduce the CFG data. On average 11 of the actual top-20 proteins were in the model’s top-20 predictions (95% CI = 6–15). For reference, selecting 20 of the 1257 sample at random will only match 11 of top 20 samples 10^−55^% of the time (see Supplementary information for calculations). The top-20 predictions included the protein with the highest RFU value for 78% of the glycans. Full lists of the top-20 predictions are included in Supplementary Table S9. A weak correlation (Pearson product-moment correlation coefficient: –0.4) between the number of entries common to both top-20 lists and the MSE of the glycan indicates that even in samples with poor MSE (e.g., Fig. 5D) can effectively predict the top-20 glycans.

**Fig. 5.**
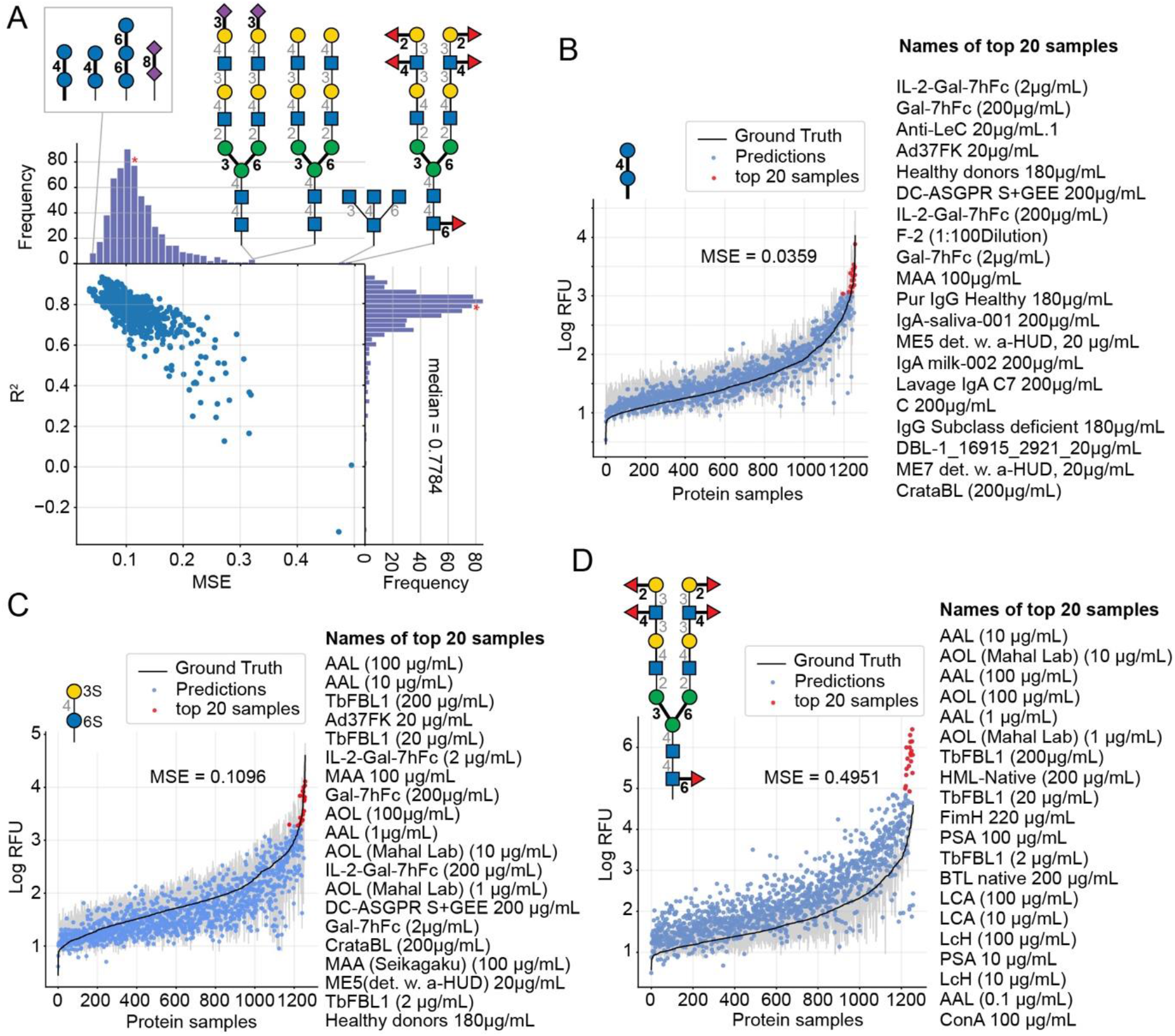
GlyNet predictions across the 1257 proteins for example glycan inputs. A plot of MSE vs. R^2^ value for 599 GlyNet predictions that describe binding of individual glycans to 1257 protein samples. Each dot represents MSE and R^2^ values from one predictive model. Panel A shows the structures of the glycans with the lowest MSE (best accuracy) and highest MSE (worst accuracy). (B-D) Plots three GlyNet predictions for three glycan inputs: the lowest MSE, medium MSE and highest MSE. In (B-D) blue dots are RFU values predicted by GlyNet and ground truth CFG values are plotted as a solid black line with grey 95% CI band, from 6 replicates. The responses in each plot are sorted in the order of increasing signal in the ground truth dataset. Red dots highlight the top 20 predicted protein samples with the highest log-RFUs. These predictions are listed (highest to lowest) on the right of each scatter plot. The glycans are B) *Glc(α1–4)Glc(α–Sp8)* (https://youtu.be/biWNApZHMP8) C) *(3S)Gal(β1– 4)[Fuc(α1–3)](6S)Glc(–Sp0* (https://youtu.be/biWNApZHMP8?t=303) and D) *Fuc(α1–2)Gal(β1– 3)[Fuc(α1–4)]GlcNAc(β1–2)Man(α1–6)[Fuc(α1–2)Gal(β1–3)[Fuc(α1–4)]GlcNAc(β1–2)Man(α1–3)]Man(β1–4)GlcNAc(β1–4)[Fuc(α1–6)]GlcNAc(β1–4)[Fuc(α1–6)]GlcNAc(β–Sp19* (https://youtu.be/biWNApZHMP8?t=598).

For further evaluation, plots of the predictions for all 599 CFG glycan instances are available in the SI. These are merged in an animation (https://youtu.be/biWNApZHMP8) stepping through the glycans (one per second) by ascending MSE of the glycan over the 1257 outputs. Specific glycans can be accessed by time (see legend of Fig. 5, Supplementary Table S9 and the legend of the YouTube video).

### Evaluation of the Glycan Binding Model by Protein Sample

Another way to assess the predictions made by GlyNet is to choose a single output from the 1257 protein-concentration outputs and study it over all 599 glycan inputs. As in the previous discussion, all 1257 predictions are available in several formats: (i) as numbers in Supplementary Table S9; (ii) as plots, two examples of which are shown in Fig. 6; and (iii) as a video (https://youtu.be/oHaFF4A22D8) indexed by protein sample in Supplementary Table S9.

**Fig. 6.**
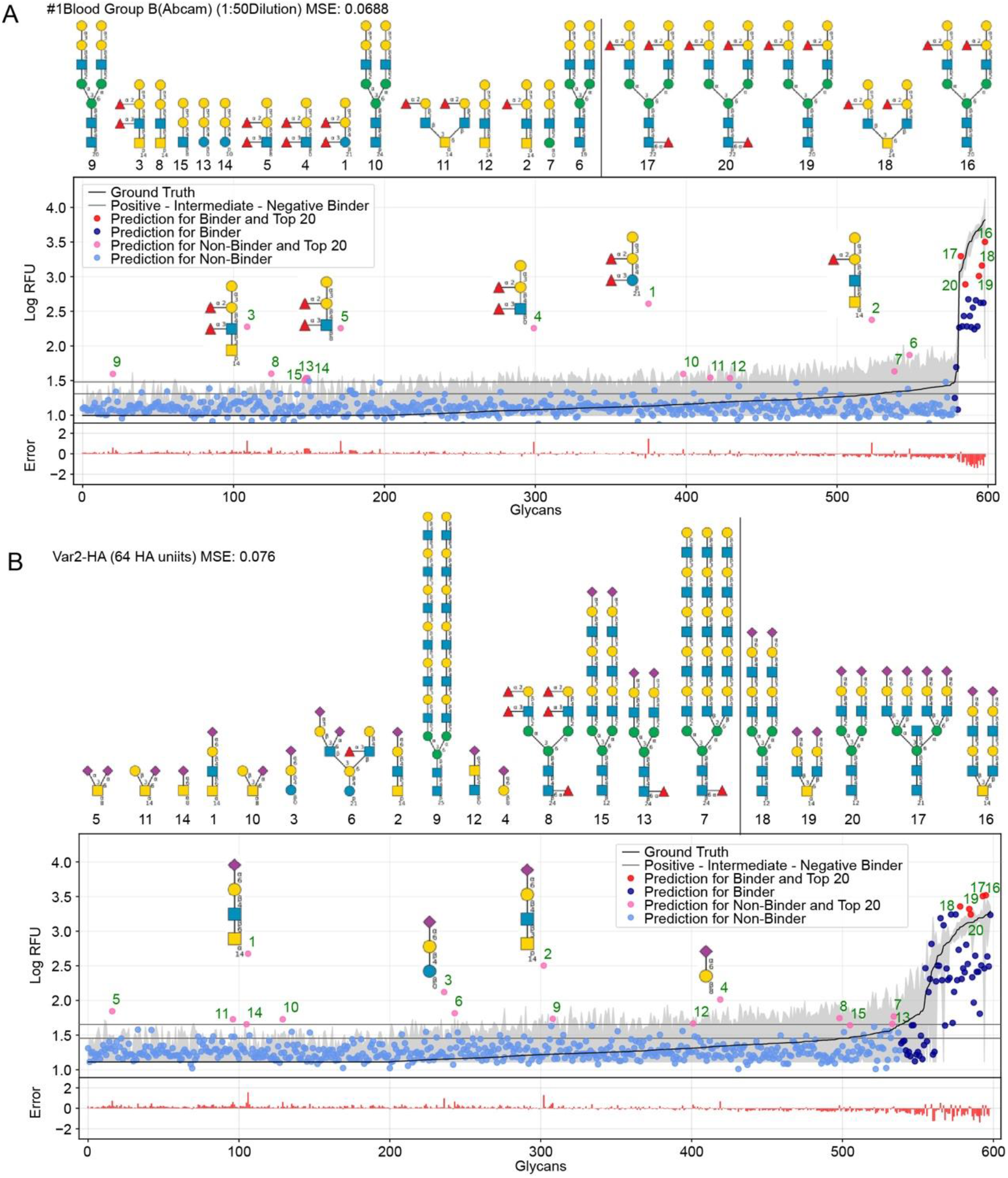
GlyNet predictions across the 599 glycans on two protein samples with well understand recognition profiles. The false positives in GlyNet’s learned outputs contain the iconic motif of the true positives. The glycans with the strongest predicted RFUs, but negative binders by the CFG data are shown with pink dots, and their structures are shown above the plots (left side). Those with the strongest RFUs and positive binders in the CFG data are red and the glycan structures are in the upper right. Also shown is the mean CFG data (black line) and its 95% CI (grey band) from N = 6 replicates. The numbers underneath the glycans represent the linkers as indicated in the CFG website (http://www.functionalglycomics.org/static/consortium/resources/resourcecoreh8.shtml), and further detail can be found in the Supplementary Table S6. A) True and false positives for an anti-blood group B antibody (https://youtu.be/oHaFF4A22D8?t=374) both contain blood group B trisaccharide. B) A sample of influenza hemagglutinin (https://youtu.be/oHaFF4A22D8?t=435) shows several positive predictions containing *Neu5Ac(a2-6)Gal(b1-4)*.

Fig. 6 shows the predictions focusing on two proteins that have well-understood glycan recognition profiles: anti-blood group antibody, and influenza hemagglutinin. To simplify interpretation of the data, we adopt the threshold for positive binders in Coff et al.^30^, (median absolute deviation based M-score ≥ 3.5) and treat the others (non- and intermediate-binders) as negative binders. Here we focus on false-positives from the GlyNet model. These are the glycan-protein pairs that were not detected as positive-binders in the CFG array data, but which GlyNet predicts to be. We find that many of these false-positives include known recognition motifs present in the true positives: e.g. GlyNet predicts that many structures with 1-2-linked fucose bind to the anti-blood group B antibody, and many glycans with 2-6 linked sialic acid (*Neu5Ac(a2-6)Gal(b1-4)*) are recognized by influenza hemagglutinin (Fig. 6). To show the generality of the observation, the aforementioned YouTube animation steps through all 1257 outputs (by ascending MSE) and true- and false-positives are shown with red and pink dots respectively. In these, we found that many of the false-positive predictions made by GlyNet have structural similarities with accepted true-positives.

GlyNet with 1257 outputs produces a total of 1257 prediction akin to those described in Fig. 6; each can be characterized with its own MSE and R^2^ value. The population median of MSE of these predictions is 0.099 (lower than MSE=0.11 in Fig. 5A) but we note two major difference: the MSE for individual samples have a wide dynamic range from 0.007 for a peanut agglutinin sample (PNA 1 μg/mL) to 0.624 for a bacterial sample (HA70 complex-Alexa 500 μg/mL). Unlike models that predict binding of 1257 protein samples to one glycan, models that predict binding of 599 glycans to one protein sample exhibit no obvious correlation between MSE and R^2^ (Supplementary Fig. S3). To analyse this observation, we examined the relationship between MSE, R^2^ and the number of glycans that bind to a protein sample. The latter can be defined either as (i) number of glycans with RFU unit(s) that are factor of 10 above the “background” RFU (Supplementary Fig. S4A) or as (ii) number of glycans above the threshold for positive binders based on median absolute deviation (or M-score^30^, Supplementary Fig. S4B,C).

When a protein sample bound to no glycans, GlyNet yielded the lowest MSEs (Supplementary Fig. S5). While accurate, this outcome is not interesting: the model learns that the output is low for every input glycan. It yields a low-signal near-random output with only small differences from the reference data (see Supplementary Fig. S5 or refer to the first 20 seconds of https://youtu.be/oHaFF4A22D8). The R^2^, a measure of correlation between prediction and ground truth, for these samples is low because there is no correlation between weak signal of the ground truth and the relatively random output of the model (Supplementary Fig. S5). This happens because MSE is the loss function whereas R^2^ was not evaluated or controlled during the learning process. As the number of strongly binding glycans increases, the MSE of the prediction increases (i.e., absolute error of the prediction increases); at the same time, the R^2^, also increases (relative quality of prediction also increases). The 20-50 protein samples with the highest MSE values bind to many glycans (e.g., Supplementary Fig. S3D or https://youtu.be/oHaFF4A22D8?t=1187 and onwards). They are the most chal-lenging for GlyNet to learn. Notably, the single-output models are no better on these samples, yielding similar performance (Supplementary Fig. S6). While these few samples are outliers in the CFG data, it would be interesting to find other learning architectures and/or encoding methods that can improve predictions for these samples.

### Evaluation of the novel predictions made by GlyNet Glycan Binding Model

Perhaps the most valuable use of GlyNet is to extend the CFG data to additional, untested glycans. We anticipate that this will aid future experiment design by identifying additional glycan structures that interact with a protein of interest, or given a glycan structure, proteins it is likely to interact with. As an example, we used GlyNet to estimate RFU values for 4160 additional glycans that have been previously synthesized or isolated and deposited onto GlyTouCan depository. Fig. 7 examines the predicted responses across all the additional glycans of individual proteins within the 1257 outputs. Animations scanning through all the output proteins are available (https://youtu.be/468Rj9ynDW4 alpha-betically by protein, and https://youtu.be/pa_6nO0Zl64 clustered by similar RFU patterns). The RFU predictions used in these plots and an index of timestamps by protein sample are available (Supplementary Tables S8 & S9).

**Fig. 7.**
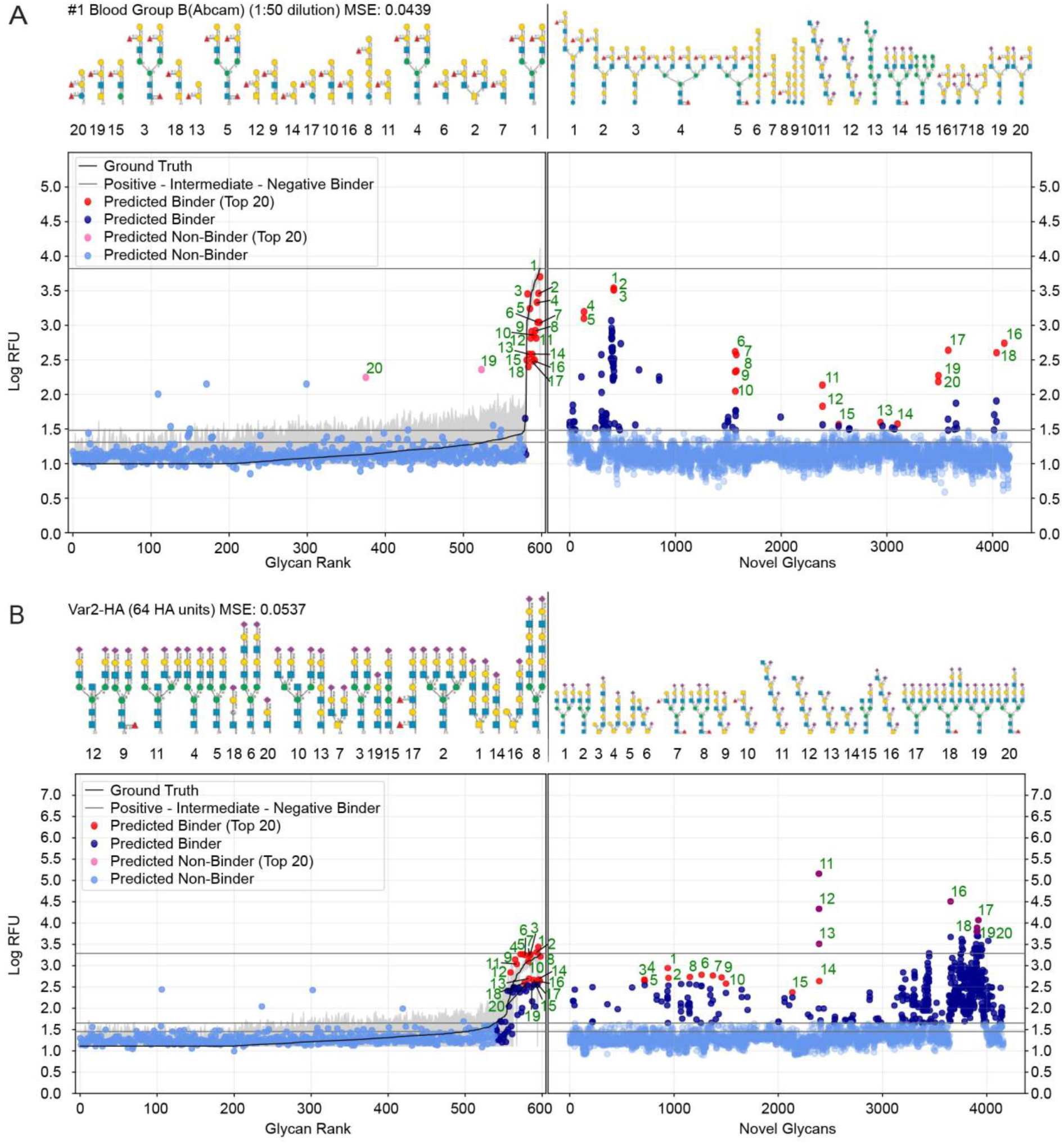
Prediction of glycan binding of Blood Group B (1:50 dilution) and Influenza Var-2-Hemagglutinin on a set of 4160 novel glycans. On the left of the plots are the GlyNet predictions across the 599 CFG glycans and the 20 glycans with the highest log-RFU values (strongest binding) are highlighted in red with structures shown above the plots. On the right are predictions of glycan binding on the set of 4160 novel glycans and similarly, 20 strongly binding glycans from across the set (red dots) are highlighted and their structures drawn above the plots. Also shown in the reference mean CFG data (black line) and it’s 95% CI (grey line) with N = 6. Glycan structures drawn with DrawGly-can-SNFG^46^.

We included a horizontal line on the plots marking the maximum RFU seen in the CFG data. For ~17% of the protein samples (213 of 1257 cases), at least one glycan is predicted to have a response more than one log(RFU) unit higher than that any of the recorded CFG glycans. In other words, the model predicts 10x more binding from one or more novel glycans, than from any glycan the model was trained against. Similarly, in ~ 4% of the protein samples (46 out of 1257 cases), GlyNet predicted glycans with RFU more than 100 times higher than RFU or any recorded CFG glycan (see Supplementary Table S8 for a list of all responses). We analyzed the ten glycans predicted to have the strongest binding for each output and observed that the distribution of strong binders was skewed towards a specific subset of the glycans. Of the 4160 glycans, only 750 glycans (18%) were present in any of the predicted top-10 binders, the remaining 3410 glycans never appear (list of these glycans available in Supplementary Table S9). Within the 750 glycans, the distribution was further skewed, with 50 glycans appearing in the top-10 of 46% of the samples (Supplementary Table S10). The majority of these 50 glycans belong to only 4 or 5 distinct classes of glycans (Supplementary Fig. S7): 10 are high-mannose structures, 16 are tri- and tetra-antennary N-glycan structures with N-terminal sialylation (6x a2-3 and 2x a2-6); LacDiNac (4/50) or LacNac (25/50) and 12 glycans that contained repeats of Gal(a1-3) (4/50), GalNAc(b1-4) (4/50) or 2-6-SialoLactose repeats -[Neu5Ac(a2-6)]Gal(b1-4)GlcNAc(b1-3)-(4/50). Remarkably, the last four glycans were nominated to be top-10 binders for 505 out of 1257 protein samples (45%) The privileged binding of only a few classes of glycans might be the result of natural binding preferences or it might represent bias in the training data producing skew in the GlyNet model or encoding strategy that favours these types of glycans over the others. While experimental validation of these predictions over all 4160 glycans is infeasible, experiments using glycans in this group of 50 or in a select members of 4-5 families may be a practical way to assess these predictions and gain assurance as to the general correctness of the predictions extrapolated by GlyNet.

### Comparison with other Machine Learning Models

The ideal way to compare different systems is to run each on the same input and obtain their respective predictions, then compare their outputs with scores such as AUC, R-squares and mean square errors. Here we compare GlyNet with CCARL by Coff et al.^30^ and SweetTalk by Bojar et al.^31–34^.

The CCARL tool^30^ was trained on the same CFG glycan array datasets as GlyNet but with slightly different pre-processing. The authors binned the RFU values into low, medium, and high categories and then trained and evaluated the classifier on the simplified task of low- vs. high-class predictions, where the medium group was removed, i.e., used neither in training nor in testing. To compare GlyNet to this tool, we created a classification version of GlyNet, which replaced the activation functions on the output neurons with logistic functions that were then thresholded at their midpoint, ½. We tested this modified GlyNet on the 20 proteins from CFG given by CCARL using the same 5-fold cross-validation sets as CCARL. GlyNet yielded an AUC of 0.912, which was higher than CCARL’s AUC of 0.895 (Fig. 8A, B). GlyNet, thus, not only can achieve a better performance after the simple conversion but is also very flexible. As GlyNet is a regression model, it can be converted to perform classifications but a classification model, like CCARL, cannot be simply converted into a regression model. Given that CCARL outperforms GLYMMR^47^, Glycan Miner^48^, and MotifFinder^49^, by transitivity GlyNet outperforms these tools too.

**Fig 8.**
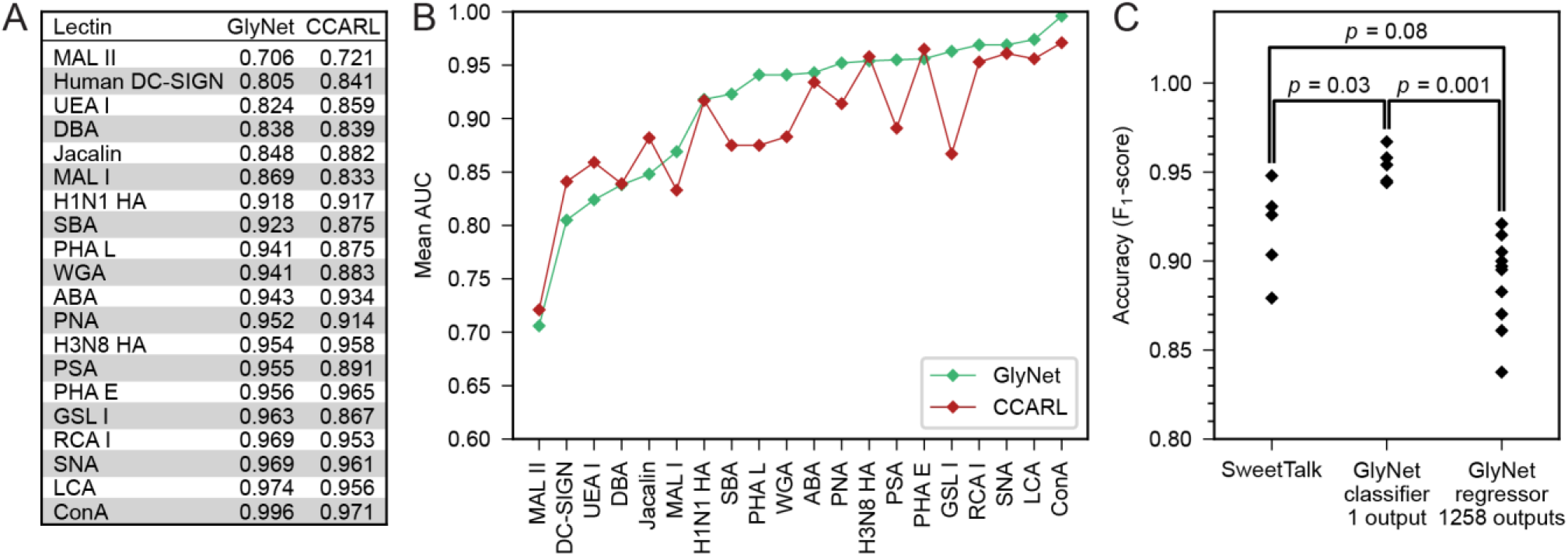
Comparison of GlyNet with CCARL and Sweettalk. A, B) Performance of CCARL and GlyNet evaluated using Area Under the ROC Curve (AUC) on the CV-folds of 20 proteins; while CCARL obtained a mean AUC value of 0.895, GlyNet obtained a better mean value of 0.912 as well as outperforming CCARL on 13 of the 20 examples. C) Comparison with SweetTalk on the immunogenicity data. Individual points are F_1_-scores for the different hold-out folds. Reported *p*-values produced by comparing F_1_-scores for individual folds by Mann-Whitney U-test (SweetTalk and 1 output *n* = 5; 1258 outputs *n* = 10; ST vs. 1 output: U = 2, common language effect size *f* = 0.08; ST vs. 1258 outputs: U = 10, *f* = 0.2; 1 output vs. 1258 outputs: U = 0, *f* = 0).

A possible explanation for GlyNet’s superior performance may originate from the features it uses. The CCLARL models used logistic regression on subgraph features after a carefully constructed set of steps chooses which features to use. In contrast, GlyNet uses the full set of substructures (up to size 3) and the neural network and the training process determines how to best use these features; an irrelevant feature will be ignored when it is assigned a small weight in the learning process. This approach avoids the risk of handicapping the final regression stage by omitting a useful feature because the earlier processing failed to identify it as important.

SweetTalk was developed by Bojar and coworkers^31–34^. Its classification of the immunogenicity of glycans is similar to GlyNet’s, as a question of whether a glycan is immunogenic can be conceptually equated to whether that glycan interacts with immune signaling proteins. To compare GlyNet and SweetTalk, we assessed classification of SweetTalk’s set of immunogenic glycans and an equal number (684) of randomly chosen human (non-immunogenic) glycans. We expanded GlyNet’s set of features in the glycan “fingerprints” to accommodate additional monosaccharides and other *q*-grams not in the CFG set. We assessed a multi-output network version which simultaneously output the 1257 RFU values and an additional immunogenic output. We found it different from SweetTalk with only weak significance (*p* = 0.08, Fig. 8C). We also tried an independent single-output classifier with a sigmoidal activation function for the output. Here, GlyNet prediction of immunogenicity has a mean accuracy (F_1_-score) of 0.954, which is a statistically significant (*p* = 0.03, Mann-Whitney U-test) increase over both SweetTalk (mean of 0.915) and the multi-output case (*μ* = 0.888, *p* = 0.001), illustrated in Fig. 8C.

GlyNet and SweetTalk use a subtly different encoding for the network input, while both decompose glycans into subunits of three monosaccharides and the two linkages between them, there are differences. For example, GlyNet also includes smaller features, allowing it to process disaccharides or even monosaccharide inputs and more importantly maintains the branched character of glycans with branched structures – i.e., we consider “V-structure” trisaccharides (Fig. 3E). To use the bidirectional LSTM of the SweetTalk system, the glycan structure is forced into a linear sequence with the end of one branch being immediately followed by the start of another. The feedforward network architecture of GlyNet offers flexibility in expanding the input features. The most important distinction is the regression (GlyNet) versus classification (SweetTalk) approaches used and a breadth of protein-glycan recognition learning tasks. It will be important in the future to assess the performance of a SweetTalk-like LSTM architecture with regression outputs on the CFG glycan array data or very similar datasets that captures a variety of protein-glycan learning tasks.

## Discussion

GlyNet is a neural network architecture trained on glycans encoded with a straightforward method that respects their graphical structure, to produce a model that can accurately predict the binding behaviors of glycans to proteins. The multi-output architecture allows the training and optimization across thousands of protein samples at once and decreases the training time which allows for a more extensive hyperparameter search. GlyNet outputs binding properties (i.e., RFUs) directly whereas many previous approaches stop at prediction of sub-motifs responsible for binding^17,18,21,22,24,26,27,29,30,48,50,51^. While it is possible to use motif information to infer binding this still requires a learning process, which is nontrivial. GlyNet is a regression model (returning a continuous range of real-valued binding strength estimates, rather than binder/non-binder classification) and we believe that a shift towards regression instead of just classification is a critical improvement in protein-glycan interaction predictions as it provides more information about the interaction allowing a deeper analysis of the outputs. Moreover, if a classifier is desired, it is straightforward to threshold the continuous output to produce classes. We believe that there is no reason to avoid building systems that predict continuous variables describing how strongly a glycan binds to specific proteins. This approach removes important decisions about choosing concentrations and binding thresholds from the machine learning pipeline and puts them into the hands of end users.

The number of possible glycans is vast, and just those of limited size containing only the most common monomers number orders of magnitude more than the number of glycans ever synthesized. The mean size of the glycans used in training is between 4 and 5 monomers. A very restricted count of possible tetra- and penta-saccharides gives 10^6^ and 10^8^ glycans, but the training set of 599 glycans covers less than 0.0001% of this space. Still, for the cases we examined, we can learn binding data given an input glycan structure. Our empirical evidence suggests that, for CFG-like glycans, we can predict the logarithmically transformed RFU values with a cross-validated MSE of 0.120. This is the first example this type of (multi-output regression) for this RFU-prediction task. As future machine learning approaches are applied to create regression models in the same data, our MSEs of 0.10 to 0.120 will serve as an important reference. Many of the 1257 protein samples, are of the same protein at different concentrations, or from different sources of nominally the same protein. A future model may be able to group such samples together to learn a dose-response trend, but we have not directly explored this.

Many of the false positives predicted by GlyNet are similar to “iconic recognition motifs;” these mis-predictions may be because our training set did not include other examples of such “near misses”. Increasing the size of the datasets, specifically inclusion of nearly identical glycans with weak protein interactions would likely improve the performance. Moreover, we know that the CFG glycan array data contains some RFU values that are inconsistent with other data sources (false negatives)^52^. Poor prediction, thus, might be a result of suboptimal training data; however, this argument can only be supported with the inclusion of additional data. Both rationales point to the need for more data and potential strategic inclusions of interactions that focus on near neighbors of the positive binders.

In this work, we used a feedforward neural network, although more complicated systems such as graph neural networks also exist. We did not use them in part because we wanted to understand the ability of this simpler system to learn and represent the protein-glycan binding for this class of data. Importantly, its results are fairly strong. Another factor motivating our preference was the relatively small amount of training data. More complex systems may require more input data to train; indeed, our empirical study suggests that even the additional complexity of the two or three hidden layer cases appears to prevent results as good as the single layer case. Note that the good results of GNNs ^53–55^ (on molecular structure graphs) were obtained on large training datasets, e.g. ~250,000 molecules from the ZINC database^56^; note that this dwarfs the 599 glycans in the CFG data we used.

As of today, few datasets are organized in machine-readable, user-accessible formats; the CFG data used in this report is a unique exception, and a rich resource for training, albeit with limits in the size and number of CFG arrays and datasets. The next generation of models will need to include other data sources and even multiple measurement techniques (e.g., glass array^4,5^ vs. bead-based array^57^ vs. frontal affinity chromatography^58^). Comparing techniques shows cross-platform variability in the generated data. To resolve this in “human learning” on protein glycan interactions, high-throughput, medium-quality data is often accompanied by high-quality thermodynamic data acquired by low-throughput techniques, such as isothermal titration calorimetry and surface plasmon resonance. We foresee three problems *en route* to next-generation ML models: (i) The data science problem of identification and organization of the needed data residing in assorted publications. Advanced techniques for mining and extraction of data (from PDFs) are needed to avoid labor-intensive and error-prone manual extraction of this data. (ii) The biochemical problem of describing of glycan presentation—for example valency, spacing, mobility, and solution vs. surface immobilized—and encoding this so that machine learning algorithms can use it. (iii) The machine learning problem of selecting effective learners that will yield useful regression models starting from a noisy inputs of diverse quality and confidence. Several other important directions are simultaneous representation of proteins and glycans (first example recently shown by Bojar et al.^34^) as well as all-atom representations of glycans with the goal of including binding affinities of glyco-mimetic compounds and non-glycan structures into training datasets. Advances in building predictive models of protein-glycan interactions requires open datasets with transparent sharing of algorithms to follow the successful path of AlphaFold2’s protein structure prediction built on PDB, CATH, psiBlast nr, and UniClust, all public datasets^59^.

## Supporting information

Supplementary Information - Includes Methods and Figures

Table S1 - Part A

Table S1 - Part B

## Data and Code Availability

A copy of the GlyNet code and the CFG derived input data is available at https://github.com/derdalab/GlyNet. The animated plots are available at: https://youtu.be/oHaFF4A22D8 (GlyNet Validation, sorted by MSE) https://youtu.be/biWNApZHMP8 (CFG glycans with respect to proteins, sorted by MSE), https://youtu.be/468Rj9ynDW4 (CFG + additional glycans, sorted by protein sample name), https://youtu.be/pa_6nO0Zl64 (CFG + additional glycans, clustered ordering).

## Acknowledgments

The authors acknowledge funding from NSERC (RGPIN-2019-04927 to R.G., and RGPIN-2016-402511 to R.D.) and NSERC Accelerator Supplement (to R.D.), GlycoNet (TP-22 to R.D.), Alberta Innovates Strategic Research Project (to R.D). Infrastructure support was provided by CFI New Leader Opportunity (to R.D.). Computation support was provided by Compute Canada. S.S. acknowledges summer research fellowship from URI and Alberta Innovates.

## Contributions

R.D. and S.S. conceived the idea. E.J.C., S.S., and N.Y. developed GlyNet and performed analysis. All authors wrote and approve the manuscript. R.G. and R.D. supervised the work.

## Supporting Information

The Supporting Information contains detail of the Methods. Fig. S1 contains a greatly enlarged version of Fig. 3a including labels and clustering dendrograms, that could not be included within the space constraints of the paper. Fig. S2 shows indistinguishable glycans with distinct structures. Figs. S3-S6 compare MSE and R^2^ to each other and to various other features of the predictions. Fig. S7 show structures for 50 frequent strong binding glycans. Table S1 contains our lists of possible small glycans. Tables S2-S6 describe the glycans and protein sample used, the pre-processed mean log-RFU values, the glycan “fingerprints,” and the glycan spacers. Tables S7, S8, S11 contain predicted RFUs used to produce Fig. 4–7 and the associated animation and Table S9 the glycan/protein time indices for the animations. Table S10 reports glycans frequently predicted to have strong protein binding.

