## Supplementary Information - Includes Methods and Figures for "GlyNet: A Multi-Task Neural Network for Predicting Protein-Glycan Interactions"

### 1. Methods

#### 1.1 Selection of CFG Glycan Array Protein Binding Data

A list of glycan array datasets was obtained from a search on the CFG website. All available CFG data downloaded for all v5.0, 5.1, 5.2 Mammalian arrays. After minor corrections of the links, in particular duplicate link removal, data was downloaded from URLs of the form: http://www.functionalglycomics.org:80/glycomics/HFileServlet?operation=downloadRawFile&fileType=DAT&sideMenu=no&objId=1006594

Where the final seven digits (the objId) varied between samples. See Supplementary Table S2 for a list of the samples and their objId numbers used.

The downloaded files are MS Excel Workbooks, and we extracted the source data from the ImaGene file formatted table. Datasets without this table were excluded from this work. The RFU for each replicate was calculated as: Mean.Signal – Mean.Background (both columns in the ImaGene table) which matches CFG’s processing described in the MS Excel files. Mean and standard deviation of these RFU values were calculated across the six replicates. This produces a wider dispersion than CFG’s numbers. Additional alignment, blank spots, and reference signals IgG, etc., were not used.

We omitted some data. Of the 611 glycans on the arrays:

i) we omitted 2 glycans not present in all three versions of the arrays,

Neu5Ac(α2-6)Gal(β1-4)GlcNAc(β1-2)Man(α1-6)[Neu5Ac(α2-6)Gal(β1-4)GlcNAc(β1-2)Man(α1-3)]Man(β1-4)GlcNAc(β1-4)GlcNAc(β-Sp13

GlcNAc(β1-2)Man(α1-6)[Neu5Ac(α2-6)Gal(β1-4)GlcNAc(β1-2)Man(α1-3)]Man(β1-4)GlcNAc(β1-4)GlcNAc(β-Sp12

ii) we omitted 1 glycan structure reported with a broken (unbalanced and therefore unparsable) branching pattern,

Gal(β1-4)GlcNAc(β1-6)[Gal(β1-4)GlcNAc(β1-2)]Man(α1-6)[GlcNAc(β1-4)]Gal(β1-4)GlcNAc(β1-4)[Gal(β1-4)GlcNAc(β1-2)]Man(α1-3)]Man(β1-4)GlcNAc(β1-4)[Fuc(α1-6)]GlcNAc(-Sp21

iii) we omitted 1 glycan structure reported to contain …Man(α1-3)[... Man(α1-3)]Man… (two substructures bound to carbon-3),

Fuc(α1-4)[Fuc(α1-2)Gal(β1-3)]GlcNAc(β1-2)Man(α1-3)[Fuc(α1-4)[Fuc(α1-2)Gal(β1-3)]GlcNAc(β1-2)Man(α1-3)]Man(β1-4)GlcNAc(β1-4)GlcNAc(β-Sp19

iv) we omitted 8 glycans contained one of five rare sugars (four or fewer instances within the CFG glycan set).

Rha(α-Sp8, GlcN(Gc)(β-Sp8, G-ol(-Sp8, MurNAc(β1-4)GlcNAc(β-Sp10

Neu5,9Ac2(α-Sp8, Neu5,9Ac2(α2-6)Gal(β1-4)GlcNAc(β-Sp8

Neu5,9Ac2(α2-3)Gal(β1-4)GlcNAc(β-Sp0, Neu5,9Ac2(α2-3)Gal(β1-3)GlcNAc(β-Sp0

We use the remaining 599 glycans and a full list of the glycans is in Supplementary Table S3. There are some minor differences in glycan name/structures found in different files, we used the latest version of the name/structure, which often corrected obvious errors in earlier versions. Our analysis ignores the linkage spacer attaching each glycan to the glass array and the stereochemistry of the stereochemistry of the anomeric position (alpha or beta) to which the linker is attached. We note that the removal of the stereochemistry of the anomeric carbon impairs the model as it cannot distinguish between cases such as Man(a1- and Man(b1- which have distinct binding properties. We also note that the information about the stereochemistry of the anomeric position is not available for 105 out of 599 glycans (see Supplementary Table S3).

Removal of linker information creates some duplicates. Specifically, in a list of 599 glycans with linkers there are 520 unique glycan structures. After removal of linker, we treat glycans as distinct entities during training although our network cannot predict different results for them. For example, (3S)Gal(β1-3)GlcNAc(β-Sp0 and (3S)Gal(β1-3)GlcNAc(β-Sp8 are treated as identical and they are encoded using the same features (Supplementary Table S5). Further details of the duplicates are available in Supplementary Table S3.

#### 1.2 Preprocessing

Initial preprocessing of the data was copied from Coff et al.^1^; constants were added to each array’s data so that the mean RFU minimum is one before transforming with log10. Then low-end noise was filtered out with minimum clamp at the 1/3rd rank position.

#### 1.3 Glycan Representation for Input to the Neural Networks

For the CFG glycan set there are 272 features, for a list see the columns of the fingerprint file (Supplementary Table S5). Features “S3”, “S4”, “S6”, and “P6” are SO_4_/PO_4_ attached to carbons 3, 4, 6, and 6 respectively. Feature names beginning with ‘[‘ are monosaccharides located in terminal positions, these are also counted within the regular monosaccharide counts. As noted in section 1.1 above, not included in the features and therefore not available to the model are details of the spacer attaching the glycan to the substrate and details of its attachment.

#### 1.4 Neural Network Architecture.

•	fully-connected feedforward neural network

•	0-3 hidden layers, all hidden layers of a common size

•	ReLU activation functions, except for linear outputs at the final stage; all with biases

•	feature count vector (fingerprint) to describe a glycan as network input

•	list of RFU values, one for each of the protein samples as output

•	implemented using PyTorch^2^

•	ADAM^3^ optimiser with weight decay and early stopping

#### 1.5 Trained model quality evaluated by 10-fold CV

Folds were created by randomly dividing (unique) fingerprints (feature vectors). Equal numbers (+/- 1) of unique fingerprints are assigned to each fold. This means that indistinguishable glycans (i.e. those with duplicate feature vectors) are assigned to the same fold. In particular, multiple instances of the same glycan distinguished by CFG with different spacers are grouped together. This prevents evaluating in hold-out a glycan case also in another fold and thus used for training. The exact number of glycans in each group varies. MSE was used to assess quality of training and CV.

#### 1.6 Parameter Optimization.

The ensure comparability when testing the effect of number of outputs on the model, the MSEs are computed over the same set of 1200 protein samples for all output sizes. A random permutation of the protein samples was produced and to evaluate the 10-output case a first model using the first ten outputs was produced, a second model from the second group of ten, until 120 models had been created. The same permutation was also used for the other sizes. The reduction to 1200 is because 1200 is highly composite, had all 1257 been used only comparable 1-, 3-, 417-, and 1257-output cases could be produced.

Using MSE and the in-fold results we tested zero through three hidden layers with 25 to 1100 neurons in each of them. By the MSE we found that a single hidden layer of 100 neurons produced an optimal MSE CV score (Fig. 4b). Similarly, we varied the ADAM weight decay parameter, while keeping 100 neurons per hidden layer and found the MSE performance was impaired as the weight-decay increased much beyond 10^-4^ (Fig. 4c). Accordingly, we fixed this parameter at 10^-4^ for subsequent work.

#### 1.7 Comparison with Other Works

For binary classification tasks, the comparisons with CCARL and the SweetTalk immunogenicity single output, the final output layer of the neural networks are changed to logistic activation functions, and the output is then thresholded at ½ to produce the binary classification.

SweetTalk immunogenicity dataset: The 684 glycans in the immunogenic_glycans_clean.csv and 684 randomly chosen human glycans from glycol_targets_species_seq.csv dataset (presumed non-immunogenic). This dataset is the same as used by the SweetTalk developers, we retain some duplicate glycans present in both groups.

This data is added as an additional “glycan array” to the CFG data by using “artificial” RFU values of 4 and 1 (after log transformation) for the immunogenic and non-immunogenic glycans respectively. Processing this also requires expanding the list of features to accommodate the additional mono-, di- and tri-saccharides not found in the 599 CFG glycan set. To convert the predicted RFU values (hold-out fold) of for this dataset to the binary classes, they were thresholding at the mean (2.5) of the two RFU input values. Changing the threshold setting from this half-way point reduced the accuracy of the results.

#### 1.8 Statistics

The RFU values obtained from CFG each have measurements are 6 replicate spots on the glycan array. We take the difference between the reported Signal Mean and Signal Background as the RFU for each spot, before calculating the mean RFU, µ, over all 6 replicates. A population standard deviation, σ, is calculated from the same six values. To estimate a 95% confidence interval the limits µ ± 1.96 σ are used, those of a Gaussian of the mean and standard deviation.

In Fig. 4 each marked data point is calculated on a different cross-validation fold. The mean and population standard deviation were calculated, and the range of the plotted error bars is the 95% confidence interval calculated with µ ± 1.96 σ, as for the RFU values. Error bars are from a single set of predictions across the 10-fold cross-validation, with a total of N = 10 points.

For comparing sets of values (see Fig. 8), Mann-Whitney U-tests are used, and two-tailed *p*-values are reported. Common language effect sizes reported are calculated by *f* = U / (*n*_1_ *n*_2_) where *n*_1_ and *n*_2_ are the sizes of the two sets being compared.

**Predictions of the top 10 or top 20 strongest binders**

Glycans from the set of 599 glycans or proteins from the set of 1257 protein samples were evaluated against a classic urn problem. For example, in Figure 5B-D, the model predicts top 20 strongest binding protein samples (from 1257 total) and it can predict a mean of 10.9 (95% CI: 6–15) of them correctly. How much better is this model than a random guess?

Given an urn with 1257 balls, 20 red and the rest white. The random model then draws 20 balls (without replacement). What is the chance that 0 of them are red? 1 of them is red? 2 are red? ... 20 are red? The answers are a hypergeometric distribution:

Pr(X = k) = K! (N - K)! n! (N-n!) / (k! (K-k)! (n-k)! (N-K-n+k)! N!)

Where N = 1257 balls total, K = 20 red balls, n = 20 draws, and k is the number of red balls drawn.

Evaluating the performance of this random model at various k:

0 from the top-20:	72%

1 from the top-20:	24%

2 from the top-20:	3.5%

…

6 from the top-20:	2x10^-5^ %	(lower edge of 95% interval)

10 from the top-20:	1x10^-12^%	(average rounded down)

11 from the top-20:	1x10^-XX^%	(average rounded up)

15 from the top-20:	1x10^-XX^%	(upper edge of 95% interval)

### 2 Supplementary Tables

#### 2.1 Supplementary Table S1 – List of Possible Small Glycans

This table lists the possible tri- to pentasaccharides we count in the introduction.

#### 2.2 Supplementary Table S2 – List of Glycans Used

This table contains details of the glycan identifiers used in the different data sources, this work, and the three versions of the CFG glycan arrays as well as the “Gene ID” used in the ImaGene style raw data tables. This is not actually a gene identifier, but the standard label used by the software which expects to analyze a DNA microarray. Entries without a number in the GlyNet column were not used in this work. See Supplementary Methods 1.1 for rationals.

#### 2.3 Supplementary Table S3 – CFG Array Information

This table lists the 1257 glycan array datasets used in this work, along with various metadata extracted from the CFG website including descriptions of the protein sample used.

#### 2.4 Supplementary Table S4 – Mean RFUs (reference from CFG)

Contains the mean RFU values after all the preprocessing steps. The table is 599 glycans x 1257 protein samples. These values are the ground truth used in learning, and used for the references for MSE calculations, and references in Figs. 4-7 and associated animations.

#### 2.5 Supplementary Table S5 – Glycan Feature Counts

This table contains are list of the glycans and the counts of the features used to describe them to the neural networks. Apart from the header, each line of the file is a different glycan and its “fingerprint”.

#### 2.6 Supplementary Table S6 – Linkers

This table contains the list of unique linkers and their chemical structure. The chemical structures of the linkers are according to the information found at the CFG website (<http://www.functionalglycomics.org/static/consortium/resources/resourcecoreh8.shtml>).

#### 2.7 Supplementary Table S7 – Predictions used in Fig. 4

This table list the GlyNet predictions that from which Fig. 4 is produced. Besides the actual predicted value, each line gives the glycan, the fold number it was assigned to, decay weight, number of hidden layers, and neurons in the hidden layers. The Repeat column contains an integer that can be used to distinguish individual runs when more than one is present.

#### 2.8 Supplementary Table S8 – Predictions used in Animations

This table contains the GlyNet output predictions used to create Figs. 5-7 and produce the associated animated plots. The reference RFU values in Table S5 are also used.

#### 2.9 Supplementary Table S9 – Timestamp Indices for Animations

This table lists times and provides links at which different protein samples and glycans occur within the animated plots.

#### 2.10 Supplementary Table S10 – Number of Top-10 Counts by Glycan

This table contains statistics about how often different glycans appear as one of the ten strongest binding glycans for each protein sample.

#### 2.11 Supplementary Table S11 – Single-output vs. Multi-output Network Predictions

Table of MSE and R^2^ values evaluation of the predictions across glycans for each protein sample from both a single-output network and a shared multi-output network.

#### 2.12 Supplementary Table S12 – Prediction for redundantly encoded glycans

This table contains summary of the predictions for a subset of glycans that have redundant encoding features.

### 3 Supporting Figures

| 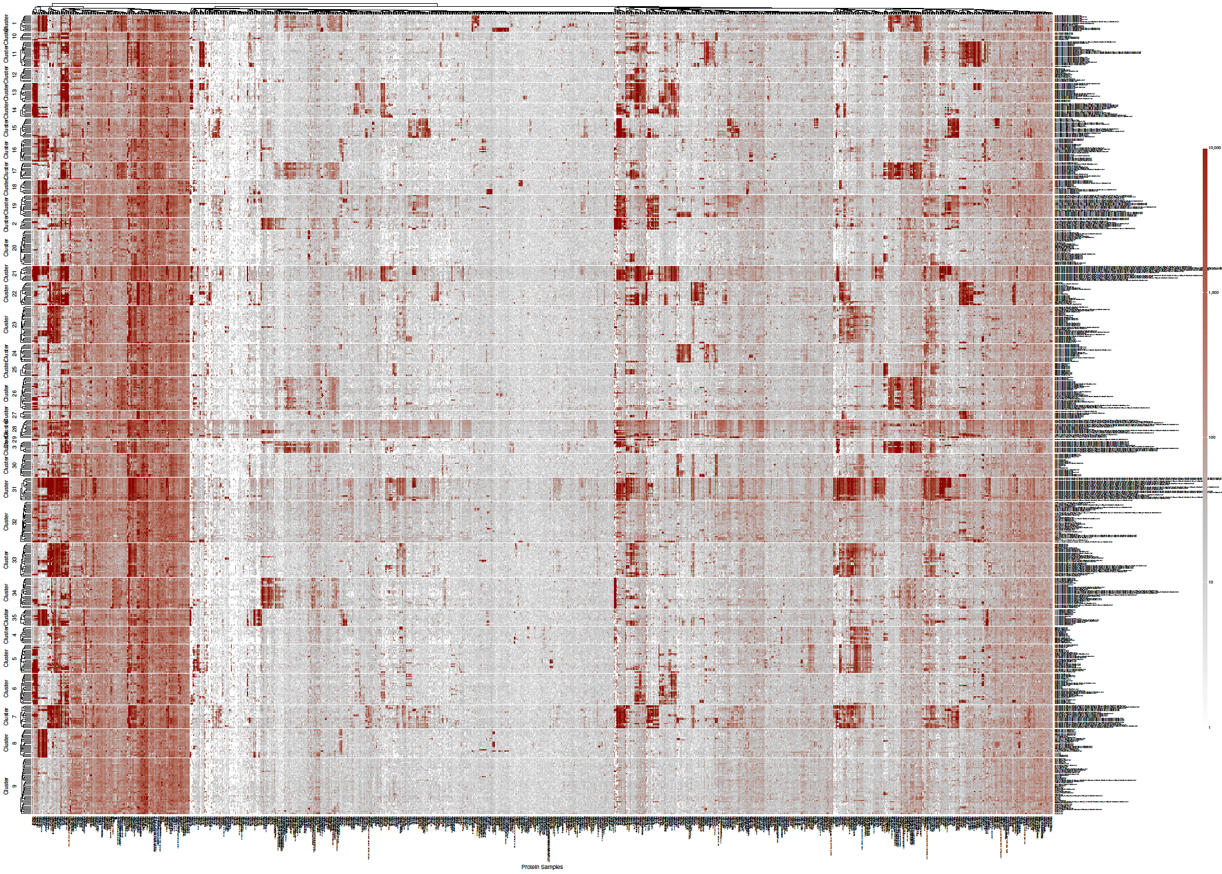 | |
| --- | --- |
| Top right corner (zoomed in)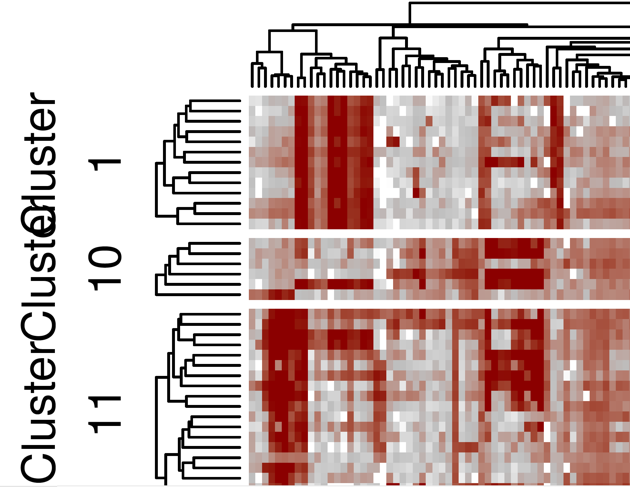 | Bottom left corner (zoomed in)  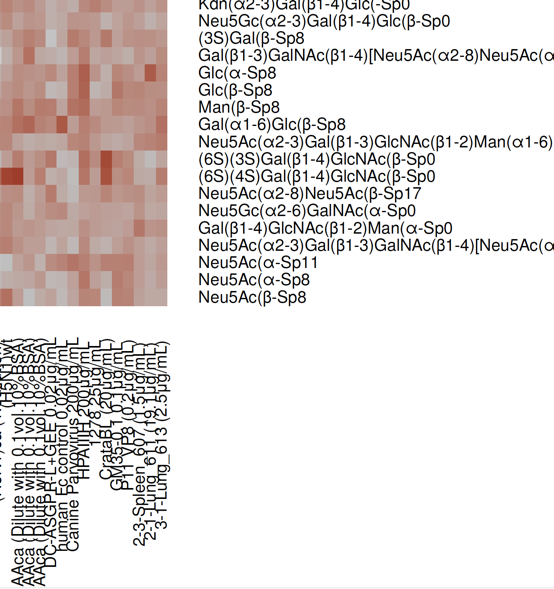 |

#### Fig. S1 – High-resolution RFU Heatmap with Dendrograms

Snapshots of the high-resolution version of the heatmap in panel Fig. 3a, this includes dendrograms showing clustering of the glycans and the glycan arrays (protein samples) as well as including textual labels for these items. Space limitations prevent these features from being included in Fig. 3a. The full figure is available in the supporting files folder as “Fig. S1 - RFU Data Heatmap with Dendrograms.pdf”

| **Isomer 1** | **Isomer 2** |
| --- | --- |
| 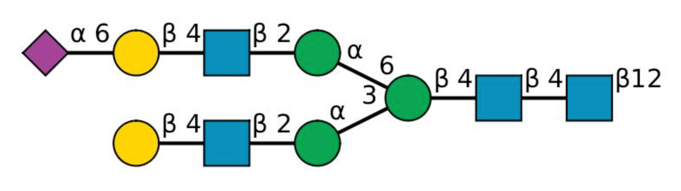  **GlyTouCan:** G75850OP  **IUPAC:** Neu5Ac(a2-6)Gal(b1-4)GlcNAc(b1-2)Man(a1-6)[Gal(b1-4)GlcNAc(b1-2)Man(a1-3)]Man(b1-4)GlcNAc(b1-4)GlcNAc(b-Sp12 | 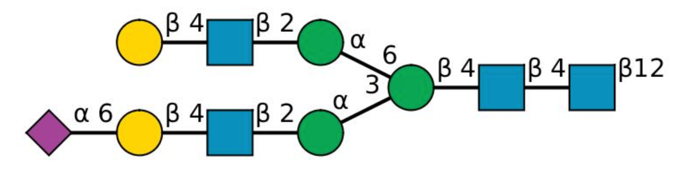  **GlyTouCan:** G91365ZQ  **IUPAC:** Gal(b1-4)GlcNAc(b1-2)Man(a1-6)[Neu5Ac(a2-6)Gal(b1-4)GlcNAc(b1-2)Man(a1-3)]Man(b1-4)GlcNAc(b1-4)GlcNAc(b-Sp12 |
| 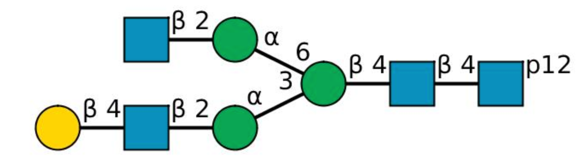  **GlyTouCan**: G99129GB  **IUPAC:** GlcNAc(b1-2)Man(a1-6)[Gal(b1-4)GlcNAc(b1-2)Man(a1-3)]Man(b1-4)GlcNAc(b1-4)GlcNAc(-Sp12 | 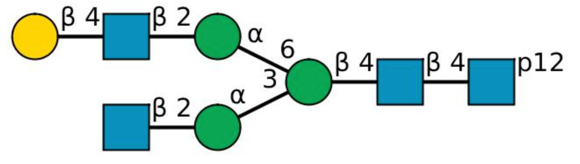  **GlyTouCan:** G86102WJ  **IUPAC:** Gal(b1-4)GlcNAc(b1-2)Man(a1-6)[GlcNAc(b1-2)Man(a1-3)]Man(b1-4)GlcNAc(b1-4)GlcNAc(-Sp12 |
| 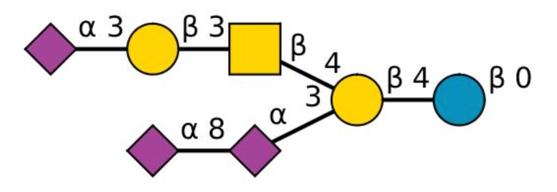  **GlyTouCan:** G40183QN  **IUPAC:** Neu5Ac(a2-3)Gal(b1-3)GalNAc(b1-4)[Neu5Ac(a2-8)Neu5Ac(a2-3)]Gal(b1-4)Glc(b-Sp0 | 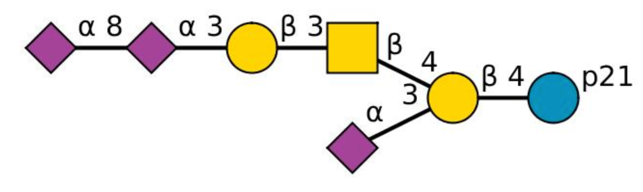  **GlyTouCan:** G97898ZO  **IUPAC:** Neu5Ac(a2-8)Neu5Ac(a2-3)Gal(b1-3)GalNAc(b1-4)[Neu5Ac(a2-3)]Gal(b1-4)Glc(-Sp21 |

#### Fig. S2 – Isomeric glycans with identical glycan feature counts

In the set of 599 glycans, we found three pairs of glycans (6 total) that are encoded by the same set of glycan feature counts. The Figure summarizes the structures, GlyTouCan codes and IUPAC names for these three pairs. Details of the encoding is available in Supplementary Table S5 - Glycan Feature Counts.xlsx

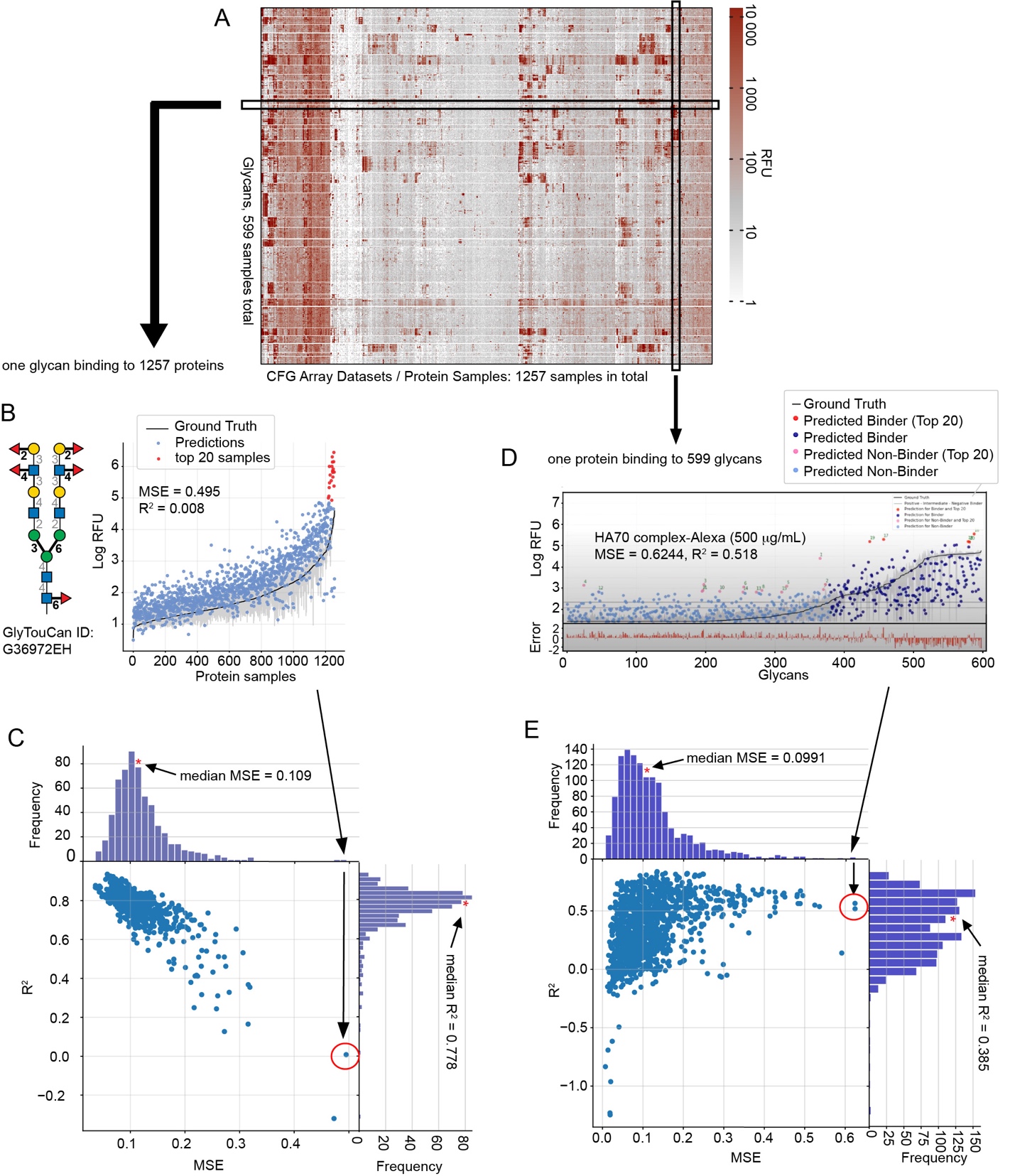

#### Fig. S3 – MSE vs. R^2^

Comparison if the two measures of quality across the glycans and the protein samples. panel (A) is copied from the Main Text Figure 3A; the vertical and horizontal slice of the heatmap represent two ways of looking at the data: (B-C) a plot describing binding of one glycan to 1257 protein samples; there are 599 plots total, all summarized as a scatter of 599 dots in MSE vs. R^2^ plot in panel (C) and available as 599-frame-long video at <https://youtu.be/biWNApZHMP8>

(D-E) a plot describing binding of one protein sample to 599 different glycans (1257 plots total). there are 1257 plots total, all summarized as a scatter of 599 dots in MSE vs. R^2^ plot in panned C) and available as 1257-frame-long video at <https://youtu.be/oHaFF4A22D8>

(B) is an example of the worst prediction of Glycan G36972EH binding to 1257 with MSE=0.495 copied from Figure 5D and the location of this prediction on the MSE vs. R^2^ plot (C) is indicated by an arrow. (D) is an example of the worst prediction for HA70 protein binding to 599 glycans and the location of this prediction on the MSE vs. R^2^ plot (D) is indicated by an arrow.

Although median MSE values are similar across (B-C) and (D-E) the median R^2^ values differ substantially. Furthermore, in (D-E) the values MSE and R^2^ are not correlated and there are numerous instances of models with low MSE (low error) but also low R^2^ (“poor fit”) and vice versa: high MSE (high error) but also high R^2^ (“good fit”). Figures S3-S5 provide further details.

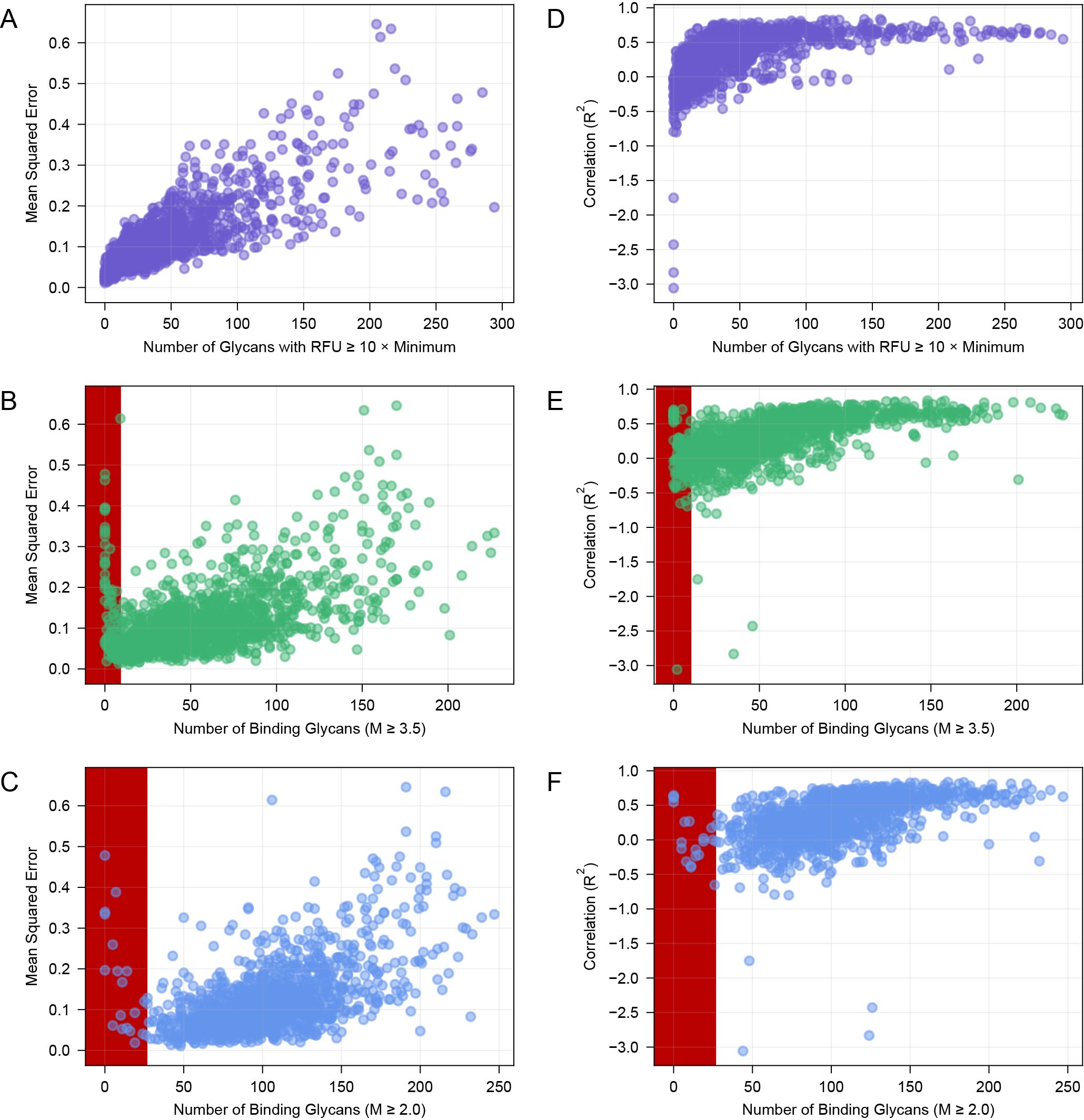

#### Fig. S4 – Correlation between MSE, R^2^ and properties of the data

(A) MSE correlates with the number of binding glycans that exhibit a response 10x above the background “N_RFU10_”. The error of the fit is low when samples has low N_RFU10_ whereas increase in the N_RFU10_ in general increases the error of the fit. (B-C) There is a weaker correlation between MSE and the number of binding glycans N_bind_; N_bind_ was defined using criteria of Coff et al.^1^. (D) in contrast to (A), the R^2^ of the fit improves with increase in N_RFU10_, whereas low R^2^ is attributed exclusively to samples with no glycans that exhibit strong response (see Figure S3). (E-F) Samples with large number of binding glycans exhibit the best R^2^ values whereas majority of the data with poor R^2^ values can be attributed to samples N_bind_ < 50. Further details are Table S9 - Timestamp Indices for Animations.xlsx

| 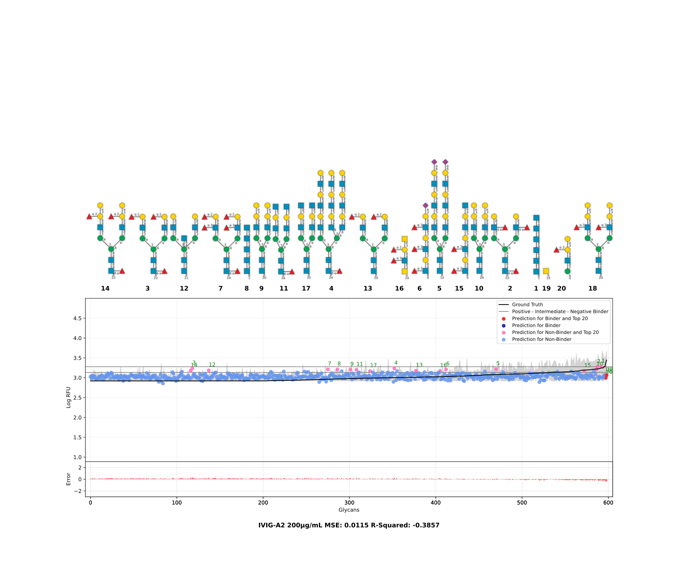 | 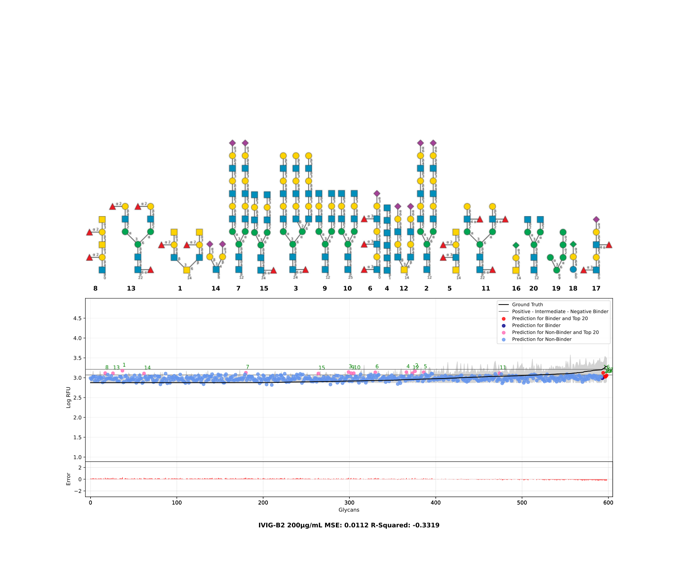 |
| --- | --- |
| 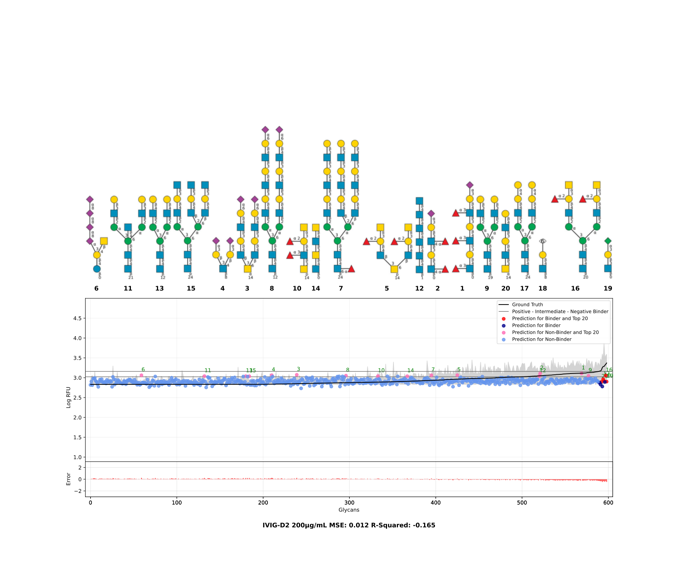 | 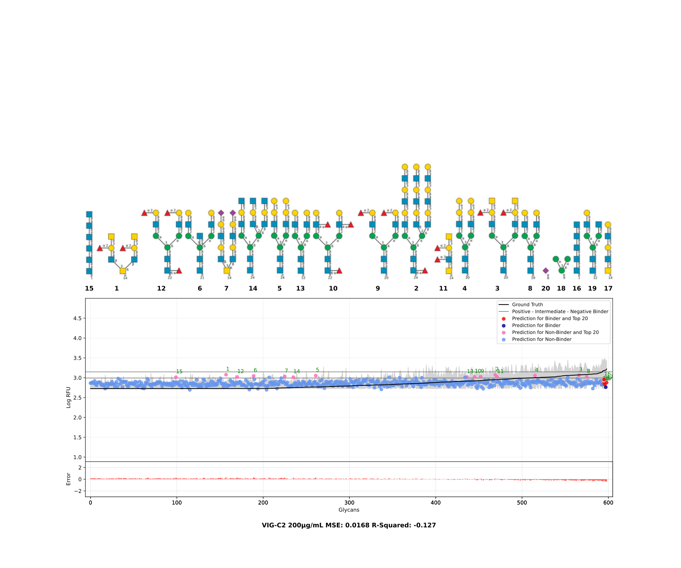 |
| 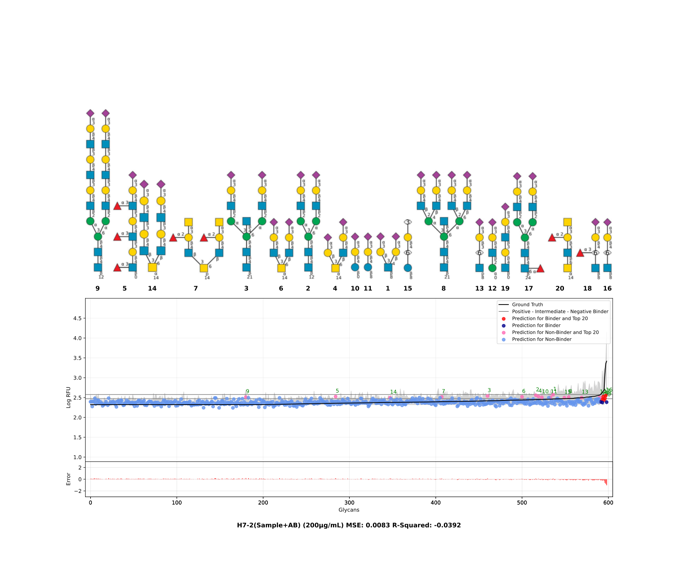 | 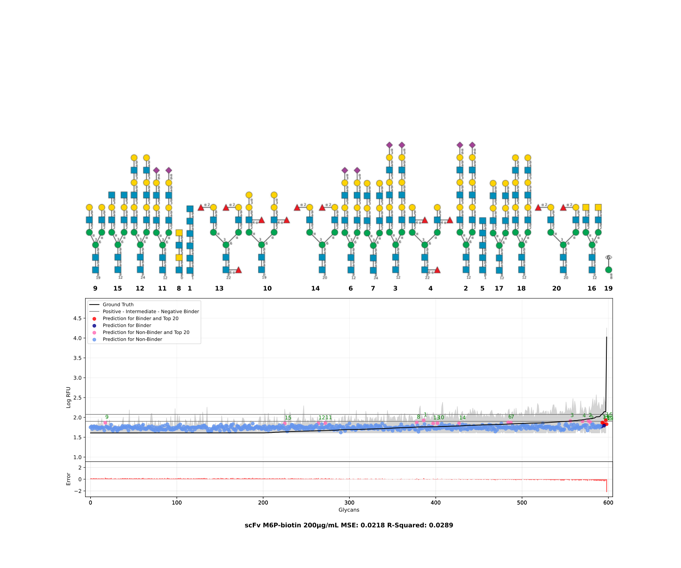 |

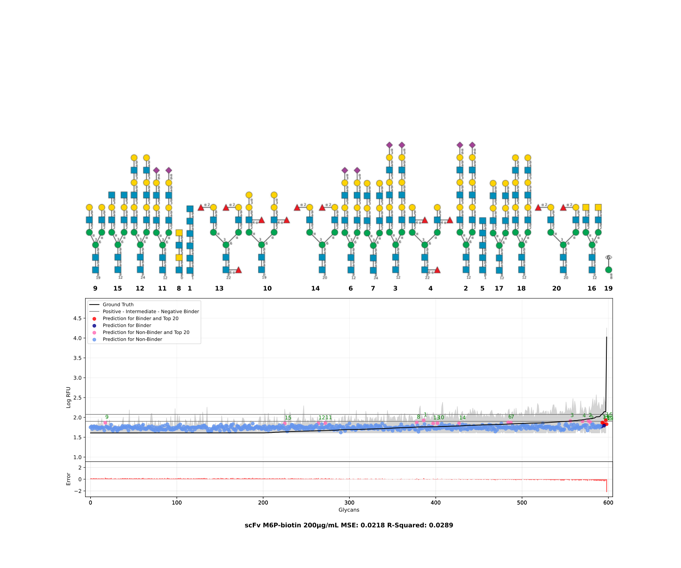

#### Fig. S5 – Examples of low-MSE-low-R^2^ data

Snapshots of six datasets that have low MSE and low R^2^. In all six cases, there are no glycans that exhibit RFU signal 10x over the baseline RFU signal and there are few or no glycans that can be classified as “binders”. Zoom in on the data in the bottom right offers some insight: the MSE is low because the absolute difference between ground truth and predicted signal is low. The R^2^ is insensitive to the absolute differences, it detects the low correlation between the ground truth and the predicted signal. Still, low R^2^ value is inconsequential to predictions in any of these cases because none of these samples contains any binding glycans. Further details can be found in Table S9 - Timestamp Indices for Animations.xlsx

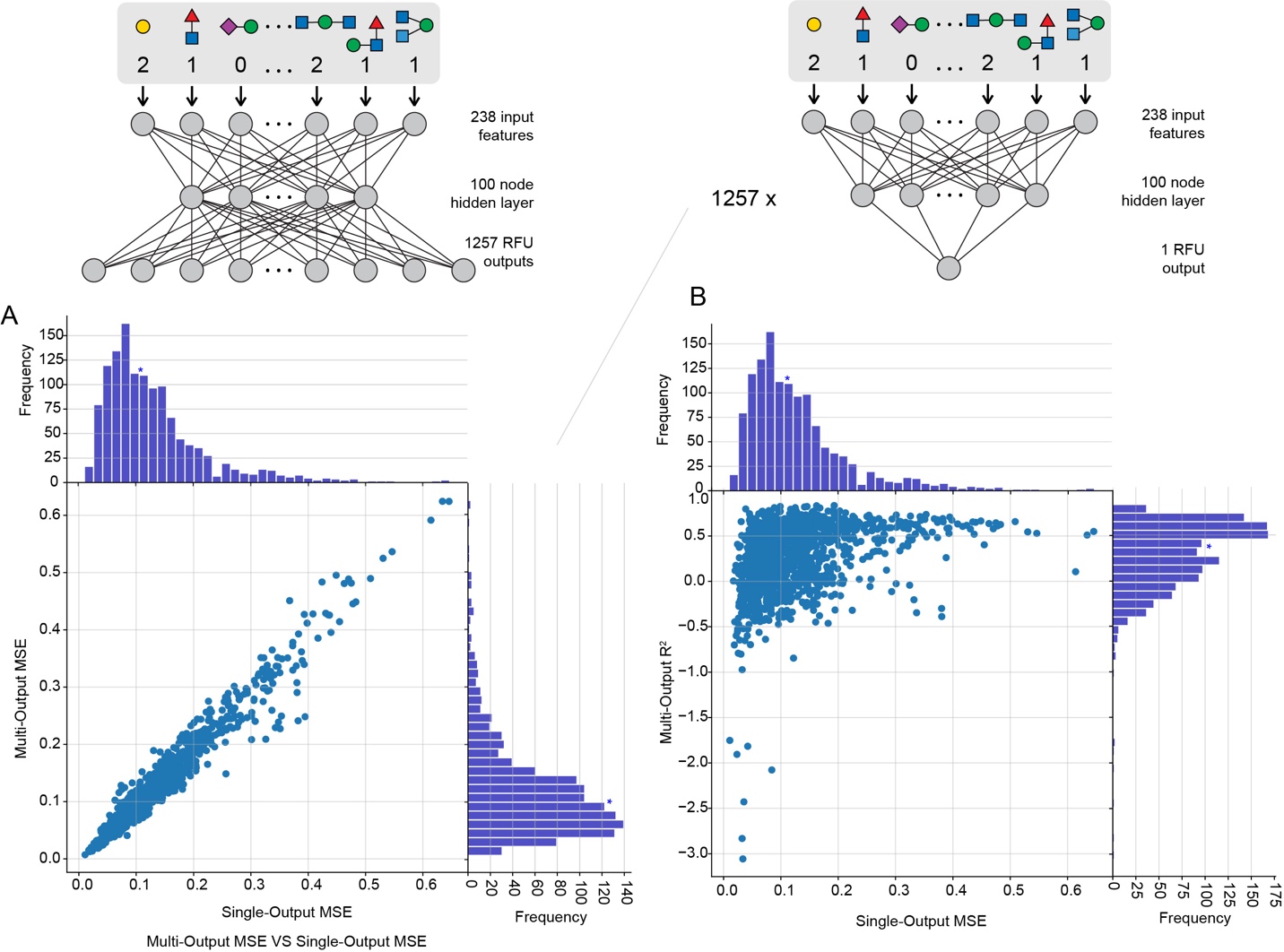

#### Fig. S6 – Correlation between MSE, R^2^ for multi-output vs single output GlyNet

(A) A scatter plot of 1257 MSE values for multi-output GlyNet vs. 1257 single output neural networks (architectures are shown on top). (B) MSE vs. R^2^ for 1257 single output neural networks: each dot is prediction for a separate protein sample. The trend of panel B is similar to the trend observed in Figure 3B (similar plot for multi-output GlyNet) but the dynamic range of R^2^ is even wider in this case. The same division of glycans into the CV folds was used for the all the single-output and the multi-output networks. We note that the switch to a multi-task network does not benefit all the protein samples, the most extreme decline in MSE being 40% (Supplementary Table S11). The decline appeared to be protein-dependent, for example a list of the negatively impacted cases contained nine samples of Ulex europaeus agglutinin (UEA) and multiple instances of a few other GBPs. This indicates that some aspects of the glycan binding properties of these GBPs was not being properly learned in the multi-output networks, and the single-output networks are able to learn these properties more effectively.

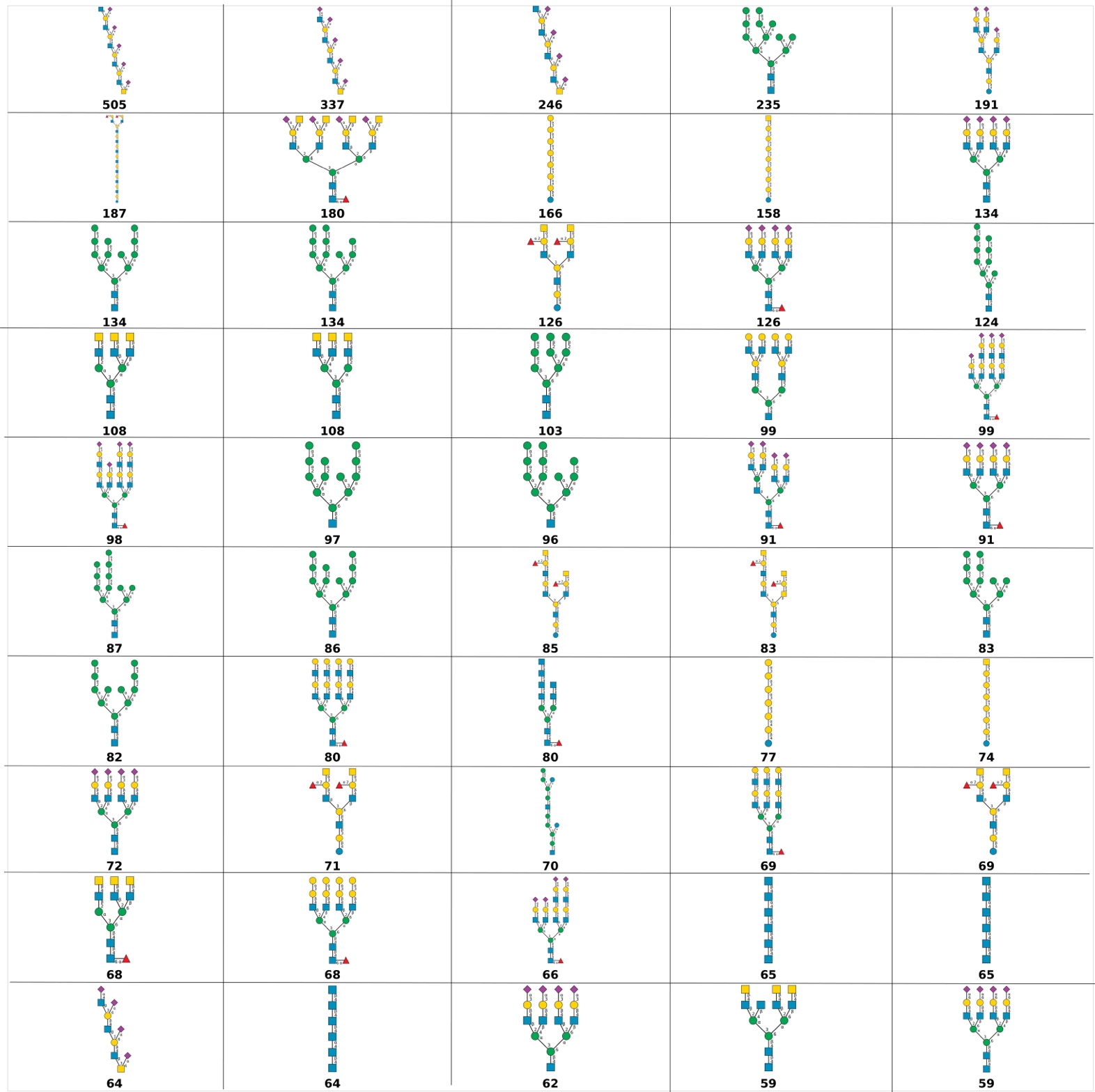

#### Fig. S7 – The Top 50 Glycans

We used GlyNet model to predict binding 4160 mammalian glycans from the GlyTouCan database to the 1257 protein samples measured using CFG glycan arrays. The image describes the top-50 privileged binding glycan structures that correspond to over 46% of the 10 strongest binders in our predictions. Numbers below the structures are the number of samples for which the glycan is a top-10 binder. The first entry bas been nominated to be a top-10 binder for 505 out of 1257 samples and the next two entries are essentially the same structure, but with fewer repeats.
